# SpatialDINO: A Self-Supervised 3D Vision Transformer that enables Segmentation and Tracking in Crowded Cellular Environments

**DOI:** 10.64898/2025.12.31.697247

**Authors:** Alex Lavaee, Arkash Jain, Gustavo Scanavachi, Jose Inacio Costa-Filho, Adam Ingemansson, Tom Kirchhausen

## Abstract

Quantitative, time-resolved 3D fluorescence microscopy can reveal complex cellular dynamics in living cells and tissues. Broader use remains limited by the difficulty of identifying, segmenting, and tracking objects of different size and shape in crowded intracellular environments in low-contrast, anisotropic, monochromatic image volumes. Objects overlap, deform, appear and disappear, and span wide ranges of size and intensity. Classical segmentation pipelines typically require high signal-to-noise data and rely on intensity heuristics with hand-tuned postprocessing that generalize poorly. Supervised deep learning methods require extensive voxel-level annotations that are costly, inconsistent across phenotypes, and rapidly become obsolete as imaging conditions change.

We introduce SpatialDINO, a fully automated self-supervised method that trains a native 3D vision transformer, based on a modified version of DINOv2 (1). SpatialDINO yields robust semantic feature maps from single channels of multi-channel microscopy that, irrespective of object shape, support object detection and segmentation directly from naïve 3D images across z-spacings and numbers of planes and different imaging modalities, without retraining or voxel annotations.

We trained SpatialDINO on a small set of confocal volumes acquired by live-cell fluorescent 3D lattice light-sheet microscopy, spanning targets of different size and shape located in crowded cellular environments, from diffraction-limited clathrin coated pits and clathrin coated vesicles to bigger structures including endosomes and lysosomes, and endosomes and lysosomes pharmacologically enlarged to highlight endosomal membrane profiles. Post-processing of the features generated by SpatialDINO allows detection and unique object identification of these objects in naïve 3D images. It also enables detection of significantly different previously unseen object classes, such as cellular plasma membranes and nuclei and even tumors in MRI scans. Finally, we illustrate its value by tracking endosomes in 3D time series, combining SpatialDINO-derived feature similarity with spatial proximity to improve association through occlusion, abrupt appearance changes, and dense packing, all conditions that have been challenging for existing methods.

SpatialDINO therefore lowers a major barrier to quantitative analysis of heterogeneous, monochromatic objects in crowded 3D cellular environments.

**SUMMARY:** Lavaee et al. have developed SpatialDINO to surmount the difficulties presented by attempting to identify, segment, and track objects in crowded volumes. SpatialDINO is a self-supervised, native 3D vision transformer trained directly on unlabeled fluorescence volumes acquired by live-cell 3D lattice light-sheet microscopy. By learning dense volumetric representations without voxel-level supervision, SpatialDINO generates features that enable fully automated detection, segmentation, and tracking of subcellular structures across a wide range of sizes and morphologies in crowded, anisotropic 3D/4D datasets acquired with different microscopy modalities. The approach generalizes across targets and imaging conditions without further training, reducing dependence on manual annotation while maintaining performance in complex cellular environments.

**SIGNIFICANCE:** SpatialDINO brings a self-supervised foundation model for analyzing 3D fluorescence microscopy images by adapting DINOv2-style joint-embedding training to learn dense volumetric features directly from unlabeled 3D datasets. By exploiting true 3D context rather than slice-wise “2.5D” aggregation, it enables automated detection and segmentation in crowded, anisotropic, low-contrast volume and enables tracking in 4D time-lapse data. SpatialDINO generalizes across targets and imaging conditions without voxel-level annotation or retraining.

## INTRODUCTION

Quantitative image-based cell biology increasingly relies on time-resolved, three-dimensional, fluorescence microscopy to relate molecular mechanism to cellular dynamics. Confocal, light-sheet, and lattice light-sheet modalities now readily yield volumetric movies over substantial time intervals, but their analysis remains rate limiting for understanding the underlying cell biology. Identifying, segmenting, and tracking objects in these data are difficult because typical volumes are low contrast, anisotropic, and diffraction limited; objects overlap, deform, appear and disappear, and span wide ranges of size and intensity in crowded intracellular environments (2).

Classical segmentation pipelines generally assume high signal-to-noise data and depend on intensity heuristics with hand-tuned postprocessing. Supervised deep learning methods can improve accuracy, but they require voxel-level annotation, which is costly to create and hard to maintain as phenotypes and imaging conditions change. These constraints motivate approaches that reduce or eliminate dependence on dense labels (2).

Recent work has begun to bring vision transformers (ViT) into bioimage analysis, with three complementary directions. First, self-supervised ViTs can learn transferable representations from unlabeled microscopy images; DINO-style model training can capture biologically meaningful variation, and yield features useful for downstream analysis without manual supervision (3, 4). Second, transformer segmentation backbones have been adapted to microscopy by fine-tuning Segment Anything and packaging interactive and automatic workflows that support microscopy annotation and, in some implementations, tracking (2). Third, transformer architectures have entered microscopy tracking more directly: Trackastra (5) learns association scores between detections within temporal windows (including dividing objects), and Cell-TRACTR (6) extends DETR/TrackFormer-style designs to end-to-end segmentation and tracking in time-lapse microscopy.

Despite this progress, two obstacles remain for routine quantitative analysis of subcellular 4D fluorescence microscopy. Most segmentation and tracking pipelines still depend on task-specific supervision or interactive prompting, and they often target “whole-cell” instances more naturally than heterogeneous subcellular structures. Moreover, many self-supervised microscopy applications operate on 2D images or “2.5D” treatments of z-stacks, failing to take advantage of native 3D context. A common workaround extends 2D foundation models to 3D by applying the encoder independently to consecutive planes and then aggregating per-slice outputs into a 3D feature volume (e.g., slice-transformer or slice-attention aggregation, averaging embeddings, or attention-based pooling) (7–9). While this strategy preserves the original 2D architecture and enables reuse of pretrained weights—avoiding the cost of training a native 3D model from scratch (7), it can limit the exploitation of native 3D context, particularly when objects overlap and move through anisotropic volumes. This limitation is not only conceptual but also practical: treating z as a weakly coupled stack discards between-slice spatial information and can compromise cross-slice consistency in downstream segmentation and tracking, an issue widely noted for 2.5D approaches (10).

To address these limitations, we have developed SpatialDINO, a fully automated self-supervised method that trains a native 3D vision transformer by adapting DINOv2-style joint-embedding training (1) to fluorescence microscopy 3D images. SpatialDINO learns dense semantic feature maps that support segmentation and tracking directly. We mitigate sparse foreground occupancy with signal-aware cropping, preserve small structures with small 3D patches, and stabilize dense volumetric feature maps by omitting explicit positional encodings, which can otherwise imprint structured artifacts.

We trained SpatialDINO on a curated set of 78 LLSM 3D time-series datasets (45,461 3 D volumes; 2.4 TB), spanning targets from small, diffraction-limited clathrin-coated pits and vesicles to larger, irregular structures such as endosomes, lysosomes, and enlarged endosomes. The resulting model has the potential to serve as a foundation model. It generalizes to structures absent from the training set, including cellular membranes and nuclei imaged by 3D fluorescence microscopy, and we have even tested it on an example of cancer-associated features in tissue MRI. It also supports robust spatial and temporal tracking of endosomes in crowded cellular environments visualized in 3D fluorescence microscopy time series. A key enabling step for tracking was deriving from the 390 SpatialDINO feature channels, an instance segmentation that uniquely labels each object by assigning a single object ID to all voxels belonging to that 3D instance.

Our results establish SpatialDINO as a powerful new tool enabling automated detection, segmentation, and tracking of crowded, heterogeneous subcellular objects directly in naïve 3D and 4D fluorescence microscopy, without voxel-level annotation, retraining, or fine-tuning.

## RESULTS

### 3D Fluorescence microscopy datasets used for training and inference

The dataset of volumetric images used to train SpatialDINO comprised single-channel fluorescence 3D time series acquired by live-cell 3D lattice light-sheet microscopy (LLSM) from living cells (78 independent imaging experiments; 45,461 volumetric stacks, ∼2.4 TB). Each volume contained ∼100–200 optical sections acquired at ∼0.25-µm z-steps, with excitation in a single channel (488, 560, or 642 nm). Most data were acquired with a home built LLSM, some with a commercial Zeiss Lattice7; both platforms used voxel sizes of 0.1 × 0.1 × 0.25 µm. The data set included gene-edited reporters spanning diffraction-limited puncta (endocytic clathrin adaptor AP2 at clathrin-coated pits/vesicles and slightly larger endosomal/lysosomal structures labeled by fluorescent EEA1 and NPC1, as well as endosomes and lysosomes containing fluorescently labeled internalized dextran. Across consecutive time points, objects often shifted in 3D between frames; treatment with apilimod, an inhibitor of the endosomal lysosomal PIKfyve phosphoinositide kinase further contributed examples with enlarged, membrane-defined NPC1-positive late endosomes (11). Collectively, the training data covered broad ranges of intensity, signal-to-noise, object density, and shape heterogeneity, from sparse puncta to crowded volumes with often touching endosomes.

The dataset used for SpatialDINO inference comprised images that were not deconvolved and were not used for training. We acquired these data as single volumes or as volumetric time series by single-channel fluorescence live-cell 3D lattice light-sheet microscopy (LLSM). The LLSM datasets included AP2-labeled clathrin-coated pits and vesicles, endosomes and lysosomes labeled with NPC1-Halo or internalized Dextran 640, nuclear pores labelled with Nup133-eGFP (12) and Hoechst-labeled nuclei. The inference data set also included live-cell 3D spinning disc confocal microscopy images of cells expressing a chimera of the membrane protein Notch fused to mNeon at its extracellular domain (13), as well as confocal datasets of cell membranes from single cultured cells and from clusters of cells in tissue, acquired by others using distinct imaging modalities, such as conventional fluorescence confocal microscopy, spinning disc confocal microscopy, two-photon Bessel-beam light-sheet microscopy (14, 15) and oblique light-sheet microscopy (Gokul Upadhyayula and Eric Betzig, personal communication); most of the images, acquired by others, had been deconvolved. Finally, the set included an MRI image of a human breast containing a tumor (9).

### SpatialDINO, overview

SpatialDINO is a fully automated, self-supervised method that trains a native 3D Vision Transformer on unlabeled, single-channel fluorescence volumes. It builds on DINOv2 (1), which learns transferable dense embeddings by training a student network to match a momentum teacher across augmented views while jointly optimizing a masked-patch (token-level) prediction objective. We trained SpatialDINO with our set of non-deconvolved low signal to noise (SNR) live-cell volumes. Despite using a limited set of data, the trained model produced accurate semantic feature maps across diverse volumes, including image types absent from training, as described below. Map quality was independent of imaging modality: the model performed comparably on low-SNR LLSM and spinning-disk confocal data, and it produced correct maps on deconvolved, high-SNR volumes.

SpatialDINO adapts DINOv2-style training to 3D by using 3D patch embeddings and fluorescence-appropriate augmentations. Because intracellular volumes contain sparse foreground, we used a k-means cropping strategy that preferentially sampled signal-rich regions. We reduced patch size to 8 voxels to preserve small structures while keeping computation tractable, and we omitted explicit positional encodings to stabilize training and suppress grid-like artifacts in dense volumetric feature maps.

During inference, SpatialDINO generated semantic feature maps that supported detection and segmentation directly from 3D volumes spanning z-spacings and different number of planes, without retraining or voxel-level annotation. We converted these features to semantic and instance segmentations using lightweight classical operators. The resulting detections also supported tracking across 3D time-lapse sequences by combining feature similarity with spatial proximity, improving associations through occlusion, abrupt appearance changes, and dense packing. Because full-volume inference at the scale of our 3D LLSM time-series data is memory-limited, we implemented a custom chunked-attention mechanism with optimized GPU kernels to enable routine processing of large 4D movies.

### Adapting DINOv2 for training on 3D fluorescence microscopy volumes

We used self-supervised learning to learn general representations from unlabeled 3D fluorescence volumes, then applied these features to downstream segmentation and spatial and temporal tracking; in principle, they could also support object classification and denoising, but we did not use them for these tasks. Our approach builds on a native 3D adaptation of DINOv2 (1), an established self-distillation method for training vision transformers on 2D images that performs well across diverse domains, including microscopy (16). Figure 1 summarizes the architectural and training changes of SpatialDINO relative to DINOv2.

**Figure 1.**
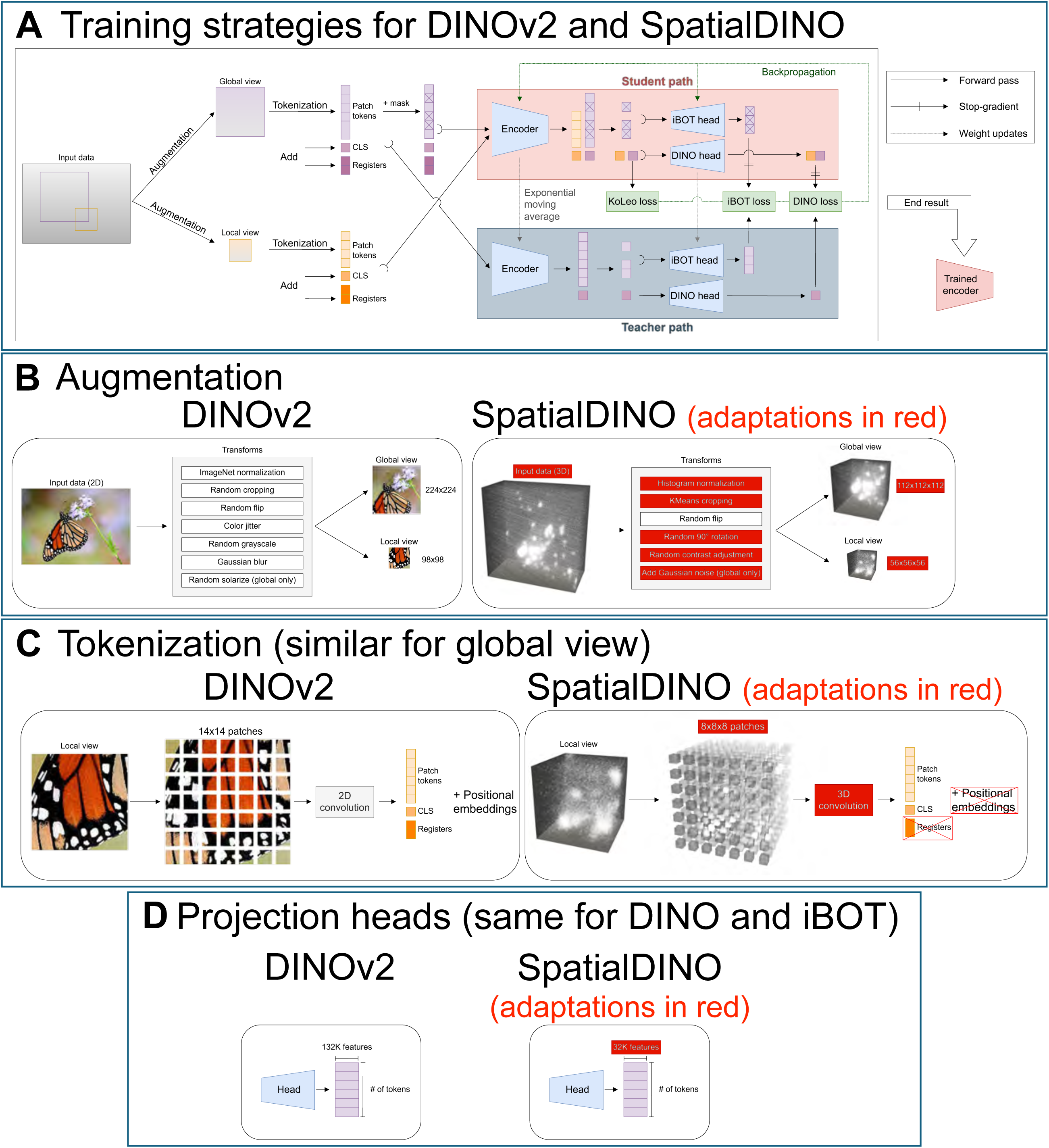
Adapting DINOv2 for 3D training. **(A)** Schematic representation of DINOv2’s training pipeline. Augmented views of a single image are passed to the network through two different paths (student and teacher). The final global and local representations from the student path are compared with the teachers via the DINO and iBOT losses, respectively, providing the learning signal for the student network, while the teacher learns “slowly” via an exponential moving average of the student’s weights. The KoLeo loss provides an extra regularization. The result of this process is a trained Vision Transformer encoder. **(B)** The augmented views in DINOv2 are 224×224 and 98×98 crops from the input 2D images. In SpatialDINO, they are 112×112×112 and 56×56×56 crops from the input 3D volumes, with a different set of augmentations highlighted in red. **(C)** The tokenization process in DINOv2 transforms 2D images into sequences of 14×14 flattened patches. To these patches we add positional embeddings to keep the spatial information between them and further include, to the end of the sequences, classification (CLS) and register tokens. SpatialDINO transforms the 3D inputs into sequences of 8×8×8 flattened patches, without positional embeddings and register tokens. **(D)** SpatialDINO’s projection heads have ¼ of the number of feature dimensions of DINOv2’s heads (32K vs 132K).

We adapted the DINOv2 architecture (Fig. 1A) to 3D inputs by replacing the 2D convolutions in the patch-embedding layer with 3D convolutions. Although similar changes have been used for 3D data that represent 2D images over time (17), this modification alone did not yield stable training on spatial 3D volumes. In pilot runs, training often collapsed after ∼10,000 iterations, producing high-norm tokens and grid-like artifacts that degraded dense feature maps.

To stabilize training and suppress artifacts, we introduced a set of changes to our SpatialDINO training pipeline (Fig. S1). The most effective change was to remove explicit positional encodings (NoPE) (18, 19). Despite the absence of positional encodings, the attention layers learned sufficient positional structure for our tasks (Fig. S1B), consistent with reports in 2D ViTs (20). To our knowledge, this strategy has not been tested systematically for 3D fluorescence volumes. Removing positional encodings mitigated the earliest collapses and enabled stable training on spatial 3D data.

We next modified the augmentation stage that generates student and teacher views for self-distillation (Fig. 1B). Because many training volumes contained sparse foreground and extensive background, standard random crops frequently sampled only background noise. We therefore replaced random cropping with k-means cropping; for each volume, we used k-means clustering to identify regions enriched in signal (k=4 for our dataset) and sampled crops preferentially from these regions (Fig. S1C; see Methods). This change reduced late-stage collapses and enabled a preprocessing option that reduced storage and training time (see Methods). Additional augmentation changes further improved robustness and modestly improved feature-map quality. We removed RGB-specific photometric transforms (e.g., color jitter, random grayscale, solarization) and replaced ImageNet normalization with histogram normalization to better handle the single channel (monochromatic) fluorescence intensity outliers. We removed Gaussian blur to preserve small, weak structures and added contrast perturbations and Gaussian noise to broaden the distribution of effective SNRs seen during training. We also added random 90° rotations to expand geometric diversity. We also adjusted global and local crop sizes (Fig. 1B) (from 224×224 and 98×98 to 112×112×112 and 56×56×56 voxels, respectively), which yielded a small improvement.

Because diffraction-limited fluorescence images often place disproportionate weight on the highest spatial frequencies within the passband, DINOv2’s default 14-pixel patch size blurred features in our volumes and disproportionately degraded small structures (e.g., clathrin coated pits and vesicles). We therefore used an 8- voxel patch (Fig. 1C and Fig. S1), the smallest size compatible with our computational constraints.

Additionally, we reduced the output dimensionality (number of prototypes) of the DINO and iBOT heads from 131k to 32k (Fig. 1D). While this change produced no obvious loss in representation quality (Fig. S1), it reduced training memory by ∼12 GB per GPU (∼30% of our usage). Our final configuration used a ViT-small backbone (1), with ∼21M learnable parameters (Table S1).

### Pipeline for SpatialDINO inference on 3D volumes

We ran SpatialDINO inference, as outlined schematically in Fig. 2, on 3D image volumes of interest. We normalized intensities with the training procedure. For time series, we estimated normalization parameters from the full movie and applied them to all frames, preserving temporal continuity and minimizing frame-to-frame flicker that could confound comparisons across time points.

**Figure 2.**
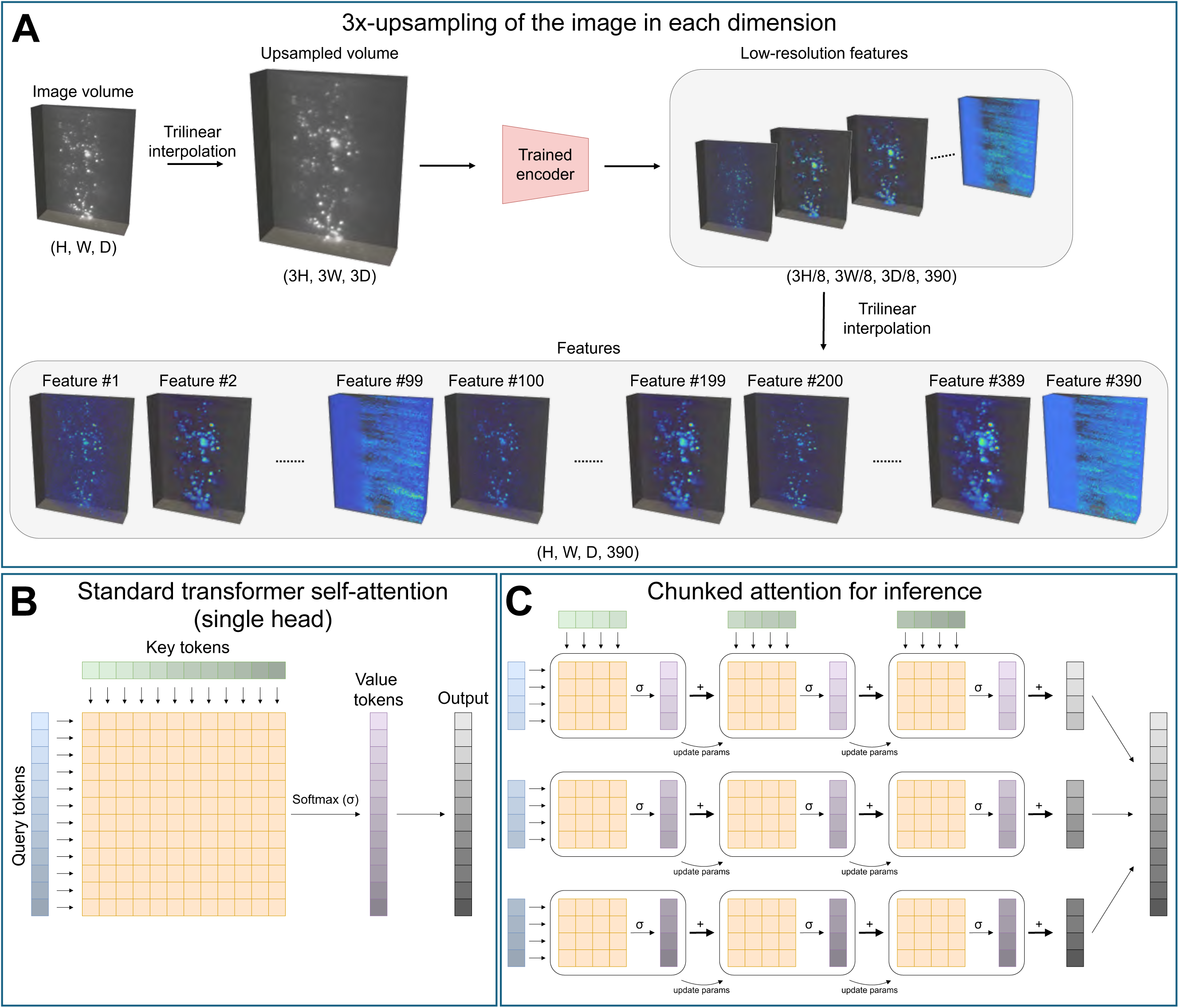
SpatialDINO inference pipeline. **(A)** Raw volumes are upsampled threefold in each dimension. For sliding-window inference, we partition the upsampled volume into overlapping blocks and process each block independently through the trained SpatialDINO encoder. We then assemble the blockwise outputs into a 3D feature map at 1/8 the upsampled resolution (3/8 of the original raw-volume resolution) with 390 feature channels, and finally upsample this map back to the raw-volume resolution. **(B)** Standard transformer self-attention (single head). The model forms a score matrix by multiplying queries by keys, applies a softmax, and multiplies the result by values to produce the attention output. The score matrix scales quadratically with sequence length and dominates attention memory and compute. **(C)** Chunked attention for inference. We avoid materializing the full score matrix by partitioning the token sequences into smaller chunks and computing attention sequentially across chunk pairs, bounding memory by the chosen chunk size while preserving full-context attention.

We upsampled each volume by a factor N (default N = 3). Because SpatialDINO outputs features at 1/8 of the input sampling, this pre-upsampling yields an effective feature sampling of N/8 relative to the raw volume, improving recovery of fine structure. When voxel spacing was anisotropic, we additionally upsampled along the coarse axis to equalize sampling. We performed all resampling with chunked trilinear interpolation to fit within GPU memory.

Because the preprocessed volumes exceeded the input size for direct ViT encoding, we modified the ViT attention module (Fig. 2B), the dominant contributor to VRAM usage, to partition each volume’s token sequence into smaller chunks and compute attention sequentially, trading parallelism for serialization. This chunked attention scheme (Fig. 2C) reduced peak memory sufficiently to run inference on standard GPUs without increasing runtime, because custom GPU kernels executed the serialized computation efficiently. (21)This yielded 390-channel feature volumes (384 patch features and 6 attention-head features) (1), sampled on a grid at N/8 of the raw resolution. We then interpolated the features back onto the original voxel grid to obtain high-resolution SpatialDINO features for downstream analyses. Figs. S2 and S3 show representative feature-map output distributions from SpatialDino applied to naïve datasets of clathrin coated pits and coated vesicles (Fig. S2) and endosomes and lysosomes (Fig. S3) not used for training. These examples illustrate the breadth, variability, and complexity of the learned representations. For each dataset, we display z-projections of all 390 feature maps from live-cell LLSM 3D images of dextran-labeled endosomes and AP2–labeled clathrin-coated pits and vesicles.

### SpatialDINO Inference Preserves 3D Object Geometry and Image Content in a PCA Feature Embedding

SpatialDINO produced dense, voxelwise feature volumes from raw 3D fluorescence datasets spanning live-cell lattice light-sheet microscopy (LLSM; not deconvolved, Fig. 4A, B), live-cell oblique light-sheet microscopy (deconvolved, Fig. 4C) and Bessel-beam/Basel light-sheet modalities (deconvolved, Fig. 4D). For each volume, we reduced the 390-dimensional feature vector at every voxel by principal-component analysis (PCA) and rendered the first three components as an image rescaled to the same 3D extent as the input.

**Figure 3.**
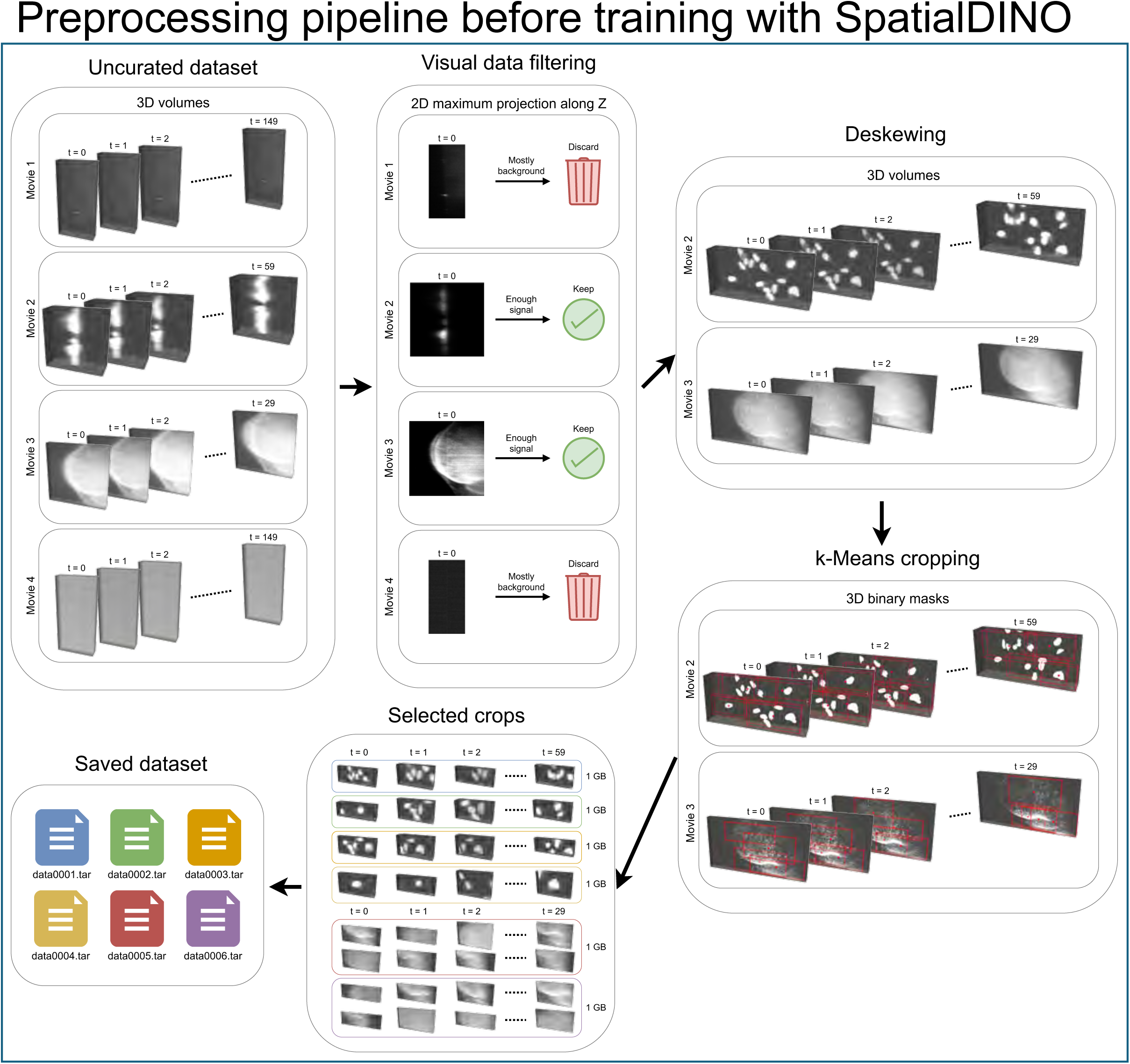
Training data preprocessing pipeline. SpatialDINO’s training data consists of 78 3D movies from various lattice light-sheet microscopy (LLSM) experiments selected from a collection acquired in the laboratory. Firstly, every movie is passed through a quick manual filtering stage where its first timepoint is max-projected along Z for visual inspection, where movies which are predominantly composed of background noise and with not enough foreground signal are discarded from the list. After this curation step, all remaining movies are deskewed to compensate the Z-dependent lateral displacement caused by our LLSM stage geometry. In sequence, we apply our k-means cropping strategy to every 3D volume in every movie, selecting k=4 signal-rich crops from each of them. These selected crops, together with their corresponding metadata, are then saved into .tar archives of 1GB of size each.

**Figure 4.**
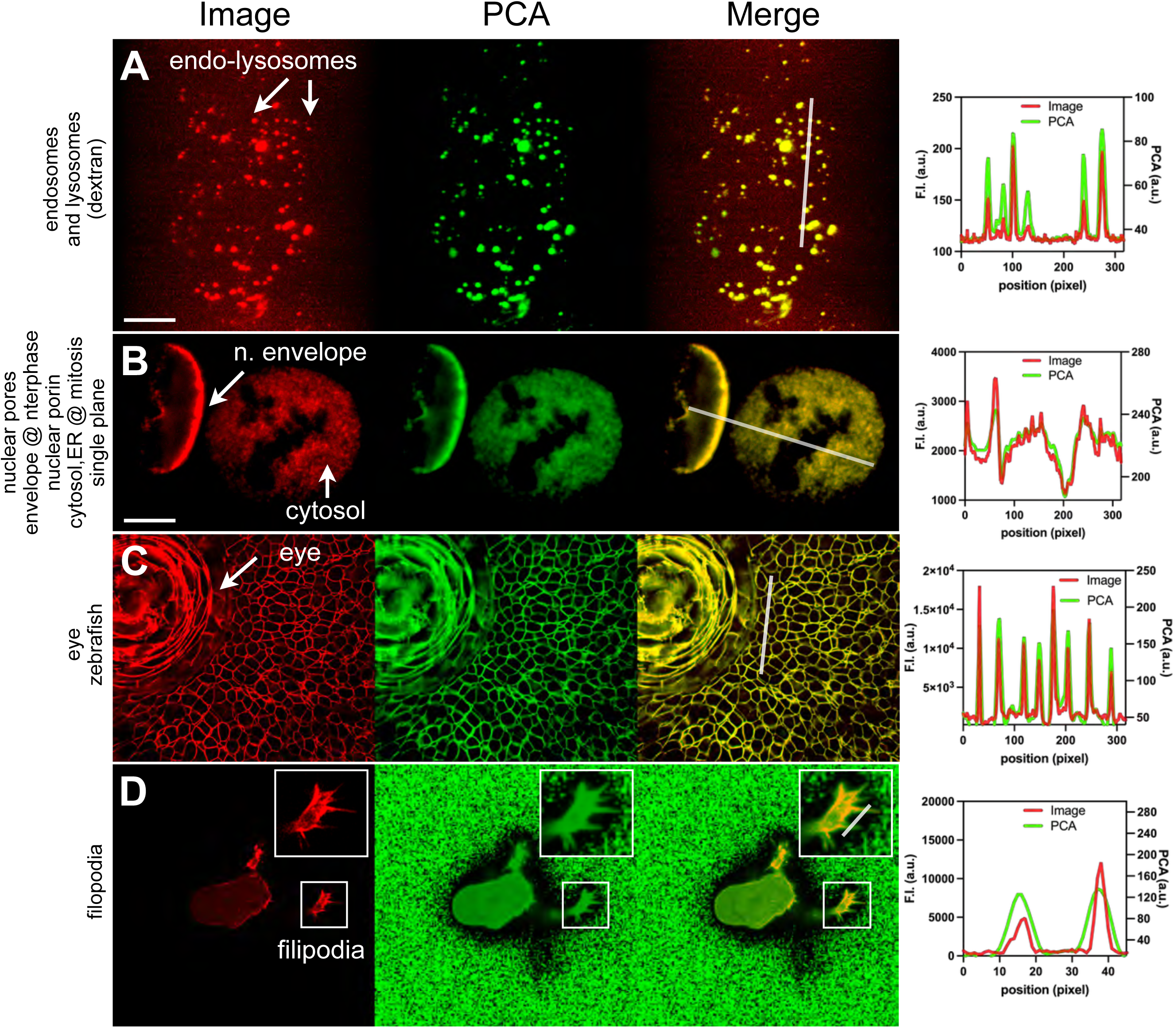
SpatialDINO inference from 3D fluorescence images yields a voxelwise feature volume whose principal components preserve object shape and image content. For each naïve example, we show (left to right) the raw image, the first PCA rendering of the SpatialDINO feature map, an overlay of raw and PCA images, and line-intensity profiles along the indicated segments. Insets mark regions of interest. PCA images report the low-dimensional embedding in image space and retain fine spatial structure. **(A)** Endosomes and lysosomes in SUM159 cells after 3 h uptake of fluorescent dextran (Dextran-Alexa647), imaged volumetrically by live-cell 3D lattice light-sheet microscopy (LLSM). Organelles appear as small 3D objects, from near–point-spread-function size to small clusters of small point-spread-functions. Similar images were included in training. Z-max projections are shown. Scale bar, 10 µm. **(B)** Halo-Nup133 in gene-edited SUM159 cells, labeled with Janelia Fluor 549 dye, imaged by live-cell 3D LLSM (acquired from (12)). This image type was not used for training. Left, nucleus of interphase cell: nuclear pores appear as spatially dense 32-mer Nup-133 puncta at the nuclear envelope, limiting optical separation. Right, mitotic cell: nuclear envelope disappears and its contents are captured by the ER; pore complexes disassemble, leaving 8-mer Nup133-containing remnants associated with the ER (12). Same single optical section is shown. Scale bar, 10 µm. **(C)** Zebrafish eye and surrounding tissue, with membranes labeled with a marker fused to mStayGold, imaged by live-cell oblique light-sheet microscopy and spatially deconvolved. We did not use this image type for training. The same single optical section is shown. Scale bar not available. **(D)** HBEC cell ectopically expressing tractin–GFP imaged by two-photon Bessel-beam light-sheet microscopy followed by spatial deconvolution (14, 15, 45). This image type was not used for training. Tractin is a transmembrane adhesion protein (46); the tractin-GFP signal concentrates at the plasma membrane and surface protrusions, with additional intracellular signal consistent with biosynthetic/trafficking pools at the endoplasmic reticulum and Golgi apparatus (14). This view using the same orientation as in Fig. 1C of (14) highlights filopodia extending from the cell surface and recognition of background ‘features’ introduced by the deconvolution procedure. Scale bar not available.

Across these naïve examples, the PCA renderings tracked the native image content and preserved object geometry: small punctate organelles, crowded nuclear-envelope puncta, extended membranes, and fine surface protrusions all appeared as coherent structures in PCA space. Overlays and line-intensity profiles showed close voxel-level correspondence between raw signal and the PCA embedding, with particular attention to object boundaries and to whether the 3D intensity distribution of each structure matched its PCA counterpart.

Because these datasets lack formal ground truth, we assessed fidelity by visual inspection at pixel resolution; in each case, the PCA representation retained fine spatial detail and respected the apparent morphology of the labeled structures. Inference for a typical 3D volume required only a few minutes. Importantly, these images were not used for training; only the endosome–lysosome example matched a training-set class, whereas the nuclear pore, zebrafish tissue membrane, and tractin–GFP datasets represented unseen image types but still yielded PCA renderings that remained spatially representative. We also compared inference on the same volumes analyzed with SpatialDINO in Fig. 4 to inference with DINOv2. For each method, we visualized the resulting feature maps by rendering the first three PCA components as RGB (Fig. S5).

SpatialDINO preserved 3D context and produced feature volumes that tracked object geometry, whereas DINOv2 features obtained by slice-wise (“2.5D”) inference showed reduced volumetric coherence.

### Foreground–Background Likelihood Mapping Converts Feature Volumes into Accurate 3D Probability Representations

Although the PCA renderings captured the gross correspondence between image content and the SpatialDINO feature volume, they often did not report object occupancy accurately. PCA emphasizes directions of maximal variance in feature space, which need not align with the spatial extent of foreground structures. For example, PCA can fail when variance is dominated by signals unrelated to the objects of interest, as in Fig. 4D, where the deconvolved background shows high variance, or in AP2 data, where clathrin-coated pits and cytosol both contribute substantial variance. In these cases, PCA largely recapitulates intensity variation in the raw data but does not separate cytosol from clathrin-coated pits and coated vesicles. We therefore implemented a simple post-processing step that converted the full 390-feature output into a voxel-wise probability map (Fig. 5). We defined foreground and background feature exemplars directly from the inferred feature volumes: for foreground, we sampled features from voxels within selected objects; for background, we sampled features from voxels in regions that clearly lacked objects.

**Figure 5.**
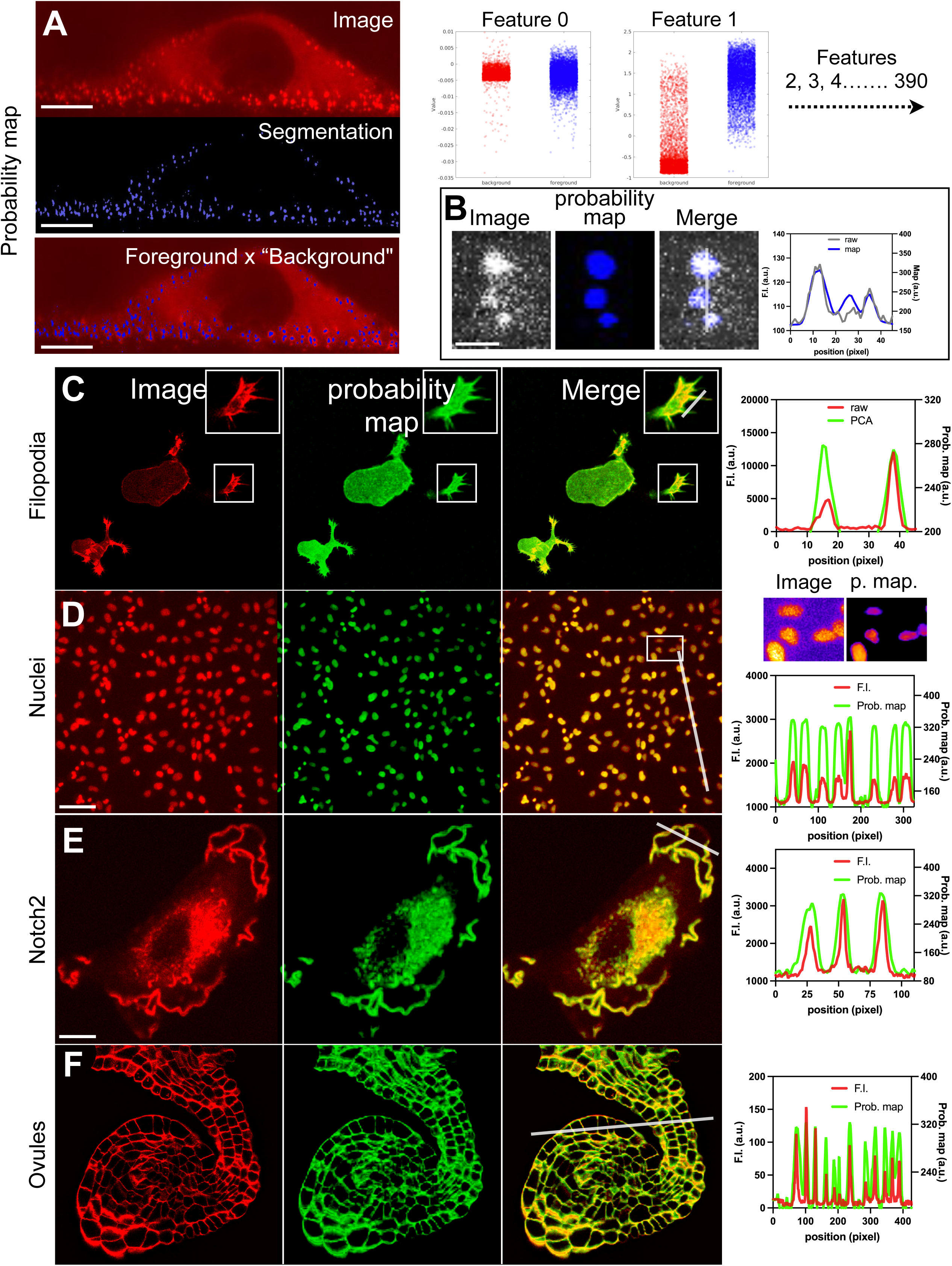
SpatialDINO inference from 3D fluorescence images yields a voxelwise probability volume that enhances representation across diverse cellular structures. Associated with Fig. S4. **(A)** Endocytic clathrin-coated pits and vesicles marked by AP2 (endocytic clathrin adaptor subunit) in gene-edited SUM159 cells expressing σ2–eGFP (21), imaged by live-cell 3D lattice light-sheet microscopy (LLSM); note that, although this is a naïve image, we used this image type for training. Coated structures are near the point-spread-function limit; diffuse background arises from soluble, unassembled cytosolic AP2. Left, top to bottom: single optical section of the raw image; fluorescence-intensity segmentation based on centroid detection using 3D CME (21) and a merge highlighting overlap used to define foreground (coated structures) and background (elsewhere). Scale bar, 10 µm. Right: distributions of voxel values for two representative SpatialDINO features (feature 0 and feature 1) for foreground and background; analogous distributions were computed for all 390 features. **(B)** Representative dextran-labeled endosome (Dextran-640; cf. Fig. 4A). Left to right: single optical section of the raw image; SpatialDINO-derived probability map; merge of raw and probability images; and line-intensity profile along the indicated segment. Scale bar, 1 µm. **(C–F)** Naïve examples not used for training. For each, we show (left to right) the raw image, the probability rendering derived from the SpatialDINO feature map, an overlay of raw and probability images, and line-intensity profiles along the indicated segments. **(C)** Single optical section of the HBEC cell expressing tractin–GFP shown in Fig. 4D. Scale bar not available. **(D)** Z–maximum-intensity projection of nuclei in cells labeled with Hoechst, imaged by spinning-disk confocal microscopy. Inset mark region of interest. Scale bar, 50 µm. **(E)** Single optical section of gene-edited SVG-A cells expressing Notch2 fused to mNeon at its extracellular domain, yielding predominant labeling of the plasma membrane (13). Scale bar, 10 µm. **(F)** Single optical section of a mouse ovule with labelled membranes from data set in (45). Scale bar not available.

We performed foreground selection using clathrin-coated pits and coated vesicles defined by local-maximum pixels detected by 3D CME (21) (Fig. 5A). For background, we sampled features from voxels in regions that clearly lacked AP2-labeled objects in LLSM volumes (Fig. 5A). We then compared the distributions of individual SpatialDINO features between the two classes and extended this scoring across all 390 features (Fig. 5A, right). From these class-conditional feature distributions, we computed for each voxel a likelihood of belonging to foreground versus background and rendered the resulting probability volume in image space (Fig. 5B, this example was obtained when applied to dextran-labeled endosomes). This probability map enhanced object representation relative to PCA, sharpening boundaries and suppressing diffuse signal, as judged by overlays and line-intensity profiles (Fig. 5B–F). We found that this approach generalized well: probability maps derived from the AP2 foreground/background definition performed robustly on other image types, including naïve examples not used for training and not used in the foreground/background definition itself. Visual inspection at pixel resolution showed that the probability renderings tracked the 3D intensity distribution of labeled structures while reducing background from soluble pools or deconvolution artifacts, yielding a more faithful volumetric depiction of cellular objects across imaging modalities and specimen types. AP2

Applying the same foreground–background likelihood pipeline (Fig. 5) to the tractin–GFP dataset emphasized the cell surface and its filopodia in a manner not achieved by PCA alone (Fig. 6). In z–maximum projections, the three-component PCA rendering preserved the main intensity patterns but remained ambiguous where diffuse intracellular signal and deconvolution-dependent background compete with thin protrusions (Fig. 6A). Converting the full 390-feature map into a voxelwise probability volume sharpened the plasma-membrane signal, enhanced filopodial continuity, and suppressed non-surface signal, yielding overlays that more cleanly separated surface structures from background (Fig. 6B).

**Figure 6.**
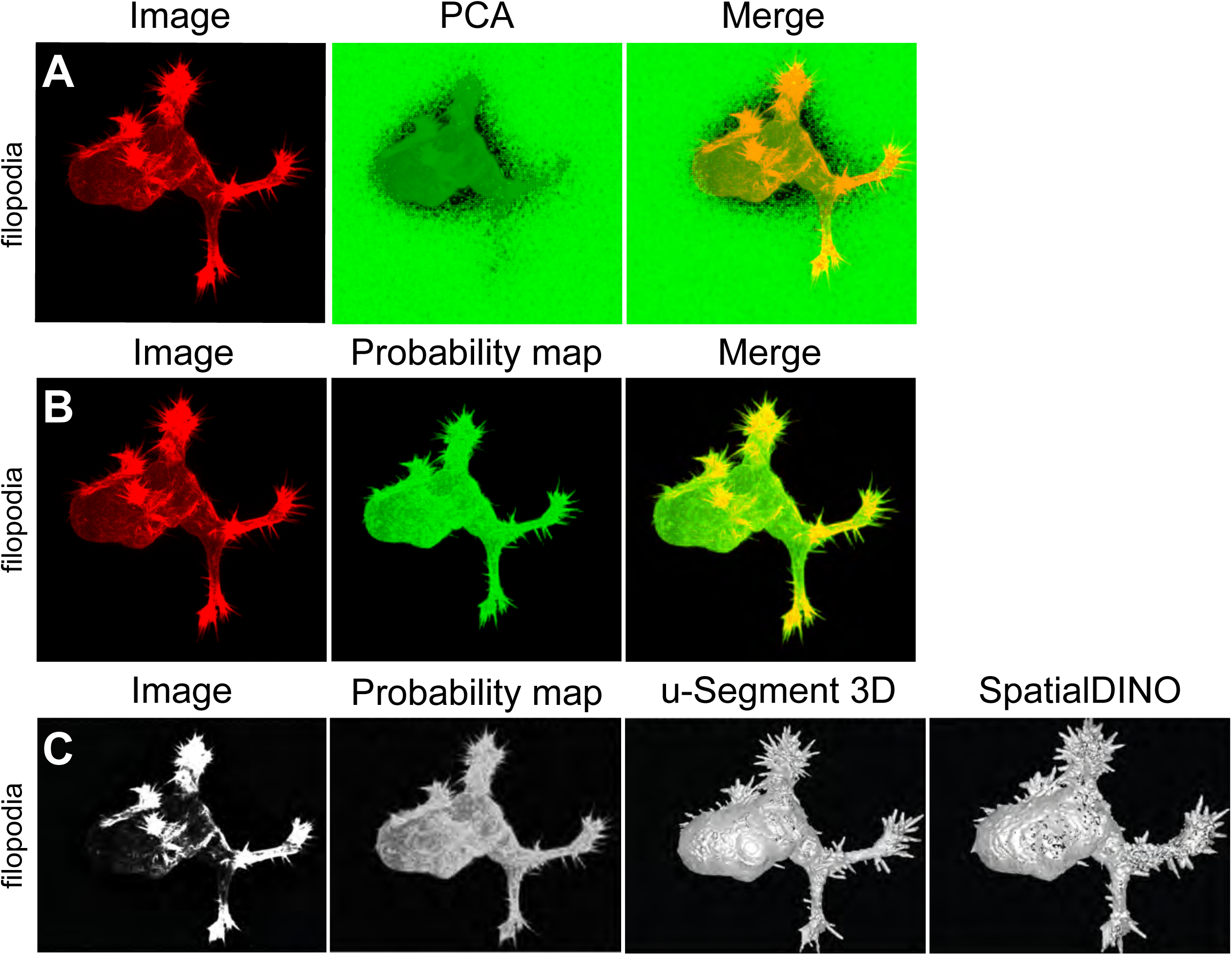
SpatialDINO inference from 3D fluorescence images yields a voxelwise probability volume that enhances representation of cell surface structures. Associated with Video 1. Same HBEC cell expressing tractin–GFP as in Fig. 4D. **(A)** Left to right: z-maximum projection of the raw deconvolved volume, the SpatialDINO-derived PCA map (from Fig. 4D), an overlay of raw and PCA images. **(B)** Left to right: z-maximum projection of the raw deconvolved volume, the SpatialDINO-derived voxelwise probability map, an overlay of raw and probability images. **(C)** Left to right: z-maximum projection of the raw deconvolved volume; the probability volume derived from SpatialDINO features; volumetric surface rendering generated with u-Segment 3D as in Fig. 1C of ref. (14); and the surface rendition of the segmentation obtained from the probability volume derived from SpatialDINO features.

We then asked whether this simple probability-volume representation recapitulates the outcome of a specialized, surface-focused workflow developed for this problem class. Using the probability volume as input, we generated a volumetric surface rendering and compared it with the u-Segment 3D reconstruction reported for tractin–GFP data (38). The resulting surface renditions were very similar, capturing the same overall cell geometry and the same population of surface protrusions (Fig. 6C). This concordance is notable because our approach relies on a generic feature-to-probability conversion applied directly to native 3D data, whereas the specialized workflow operated on 2D stacks that were converted into an inferred volume.

These examples illustrate how SpatialDINO inference combined with minimal post-processing can yield a practical route to accurate 3D representation of cell-surface structures, without tailoring the model or the reconstruction method to a specific specimen or imaging modality.

### Segmentation by PCA Signatures and Foreground–Background Probability Mapping in SpatialDINO Feature Space

We asked whether SpatialDINO features can support instance-level object identification without retraining, by using either PCA-derived signatures or the foreground–background probability maps (Fig. 7). For a controlled test, we simulated crowded 3D volumes containing endosome-like objects with heterogeneous shapes and internal intensity patterns and with low signal-to-noise ratios (SNR 3–15) (Fig. 7A, ground truth). Voxel-wise PCA values were sufficient to distinguish objects from background and co-localized with the ground truth (Fig. 7A, PCA-Seg), enabling clustering of voxels into unique, individually segmented objects (PCA-Seg-ID). The resulting per-object PCA signatures were effectively unique; we therefore assigned each segmented object a distinct pseudo color to visualize identity (PCA-Seg-ID). Across 15,004 simulated objects (∼300 objects per volume across ~50 volumes), this PCA-based approach correctly segmented 14,602 instances, yielding high precision, recall, and F1 score over the full set (Fig. 7A). We then applied the same self-supervised logic to different types of experimental data that lacked voxel-level ground truth for object identities or boundaries.

**Figure 7.**
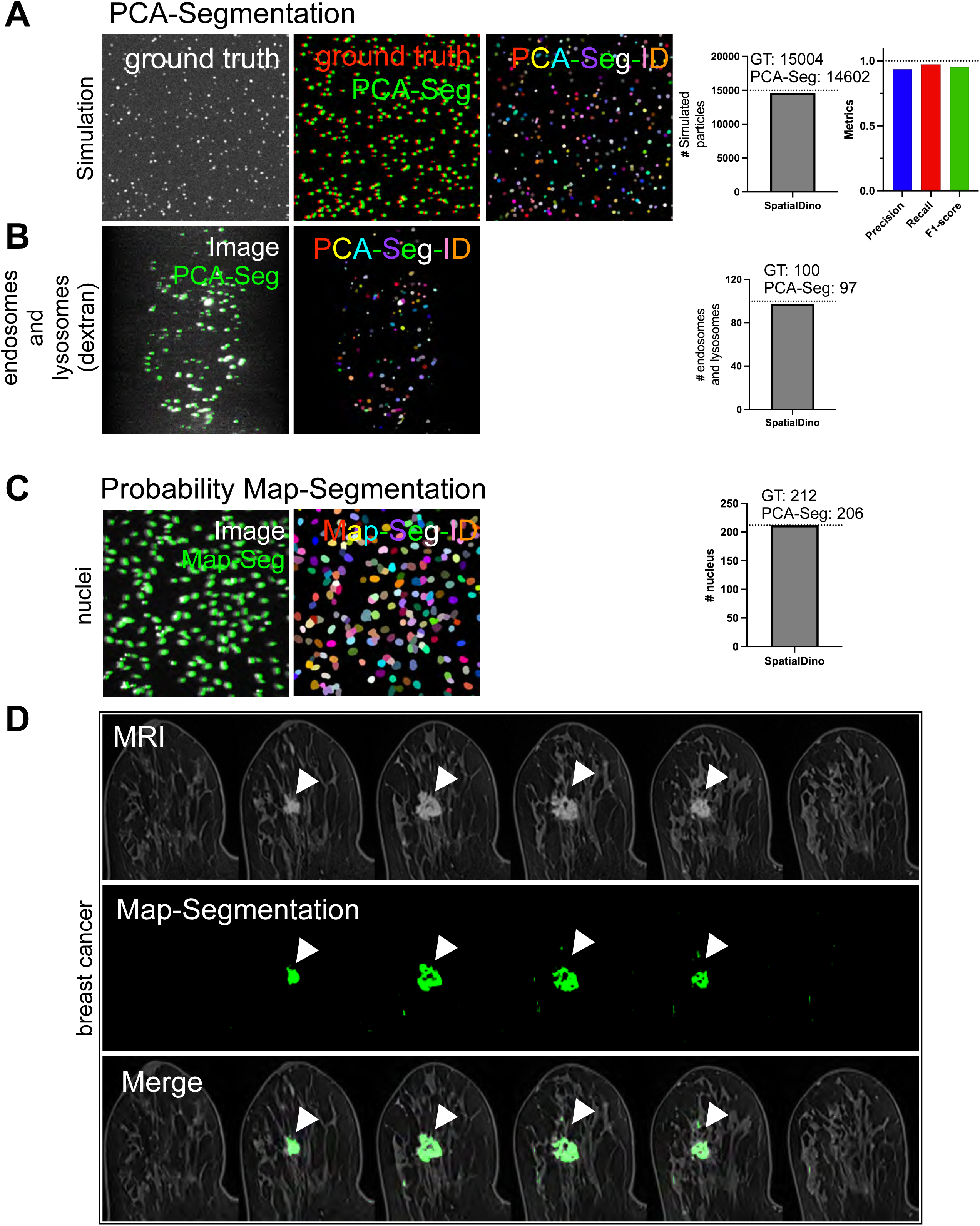
Object identification from segmentations based on SpatialDINO PCA or probability feature maps. Associated with Video 2. **(A)** Simulated endosomes. Left to right: z–maximum-intensity projection of the ground-truth objects; merge of the ground truth (red) with PCA-based segmentation from SpatialDINO (in green) shifted by 6 pixels along the x axis for clarity; and instance segmentation based on the PCA with a distinct color assigned to each object. Total number of objects (GT = 15004) and number of simulated particles correctly segmented by SpatialDINO (PCA-Seg = 14602); dotted line highlights the high precision, recall and F1-Score across all 50 time points. **(B)** Dextran-labeled endosomes and lysosomes imaged using live-cell 3D LLSM. Left to right: merge (SpatialDINO PCA-based segmentation in green) with the z–maximum-intensity projection of a live-cell 3D LLSM volume of internalized Dextran-640 (gray), shifted by 6 pixels along the x axis; and instance segmentation with a distinct color assigned to each object. Total number of endosomes and lysosomes (GT = 100) and number correctly segmented by SpatialDINO (PCA-Seg = 97). Scale bar, 10 µm. **(C)** Hoechst-labeled nuclei imaged using live-cell spinning disk confocal microscopy. Left to right: merge (SpatialDINO probability-map segmentation in green) with the z–maximum-intensity projection of a live-cell 3D spinning-disk confocal volume (gray), shifted by 6 pixels along the x axis; and instance segmentation with a distinct color assigned to each object. Total number of nuclei (GT = 212) and number correctly segmented by SpatialDINO (PCA-Seg = 206). Scale bar, 50 µm. **(D)** Breast MRI, right breast (DUKE: Breast_MRI_141 (right) from (9)). Top to bottom: axial slices showing a malignant lesion (white arrowhead), adapted from (9) (every third slice shown); probability-map segmentation from SpatialDINO applied to the full 3D MRI volume (green); and merged view. The probability map was generated using lesion foreground–background labels for the ipsilateral breast.

For dextran-labeled endosomes and lysosomes imaged by live-cell 3D LLSM, PCA-based instance assignment produced distinct identities for nearly all puncta within a volume (Fig. 7B). Of 100 organelles, 97 received unique assignments, consistent with robust object-level separability in feature space.

As a complementary path we used the probability feature map rather than PCA. For Hoechst-stained nuclei imaged by live-cell spinning-disk confocal microscopy, foreground–background likelihood mapping produced a probability volume that enabled reliable detection and instance labeling of most nuclei (Fig. 7C). We correctly segmented 206 of 212 nuclei; the missed cases corresponded to nuclei in close apposition, where touching boundaries limited separability at the image resolution.

Finally, we asked whether the same SpatialDINO trained model could be transfer and used to an imaging modality far from fluorescence microscopy. We carried this test by using SpatialDINO to infer features on a

3D breast MRI volume from a published study that had used large scale supervised training to identify malignant lesions (7). Using lesion-aligned voxels in one breast to define foreground and surrounding tissue as background, we generated a probability map from the full 390 feature volume and applied it to the paired breast. This probability-based segmentation highlighted the contralateral lesion at the location annotated by expert review in the original study (Fig. 7D). In this example, we did not restrict foreground exemplars to voxels within the cancer mass; instead, we applied a simple threshold to the probability map to capture the lesion while minimizing (but still including) unrelated structures elsewhere. Thus, a simple feature-to-probability postprocessing step can convert SpatialDINO’s volumetric embedding into a practical MRI segmentation, using only the original low-SNR 3D fluorescence volumes used as exemplars to train SpatialDINO and without imaging-modality–specific retraining.

The MRI example underscores both the generality and the current limits of our approach. SpatialDINO was trained on a limited set of fluorescence volumes, yet its inferred features were sufficient to support lesion highlighting after postprocessing. Further work will be required to tune post-inference processing so that the output emphasizes only the feature of interest, here the tumor mass.

### Feature-Based Instance Identity Enables Robust 3D Tracking Through Crowded Space

Unique instance identities within a 3D volume should enable more reliable spatiotemporal tracking than proximity alone, particularly in crowded fields where trajectories cross and nearest-neighbor assignments become ambiguous. We therefore asked whether SpatialDINO-derived object signatures can stabilize linking across time in both simulated and experimental data (Fig. 8).

**Figure 8.**
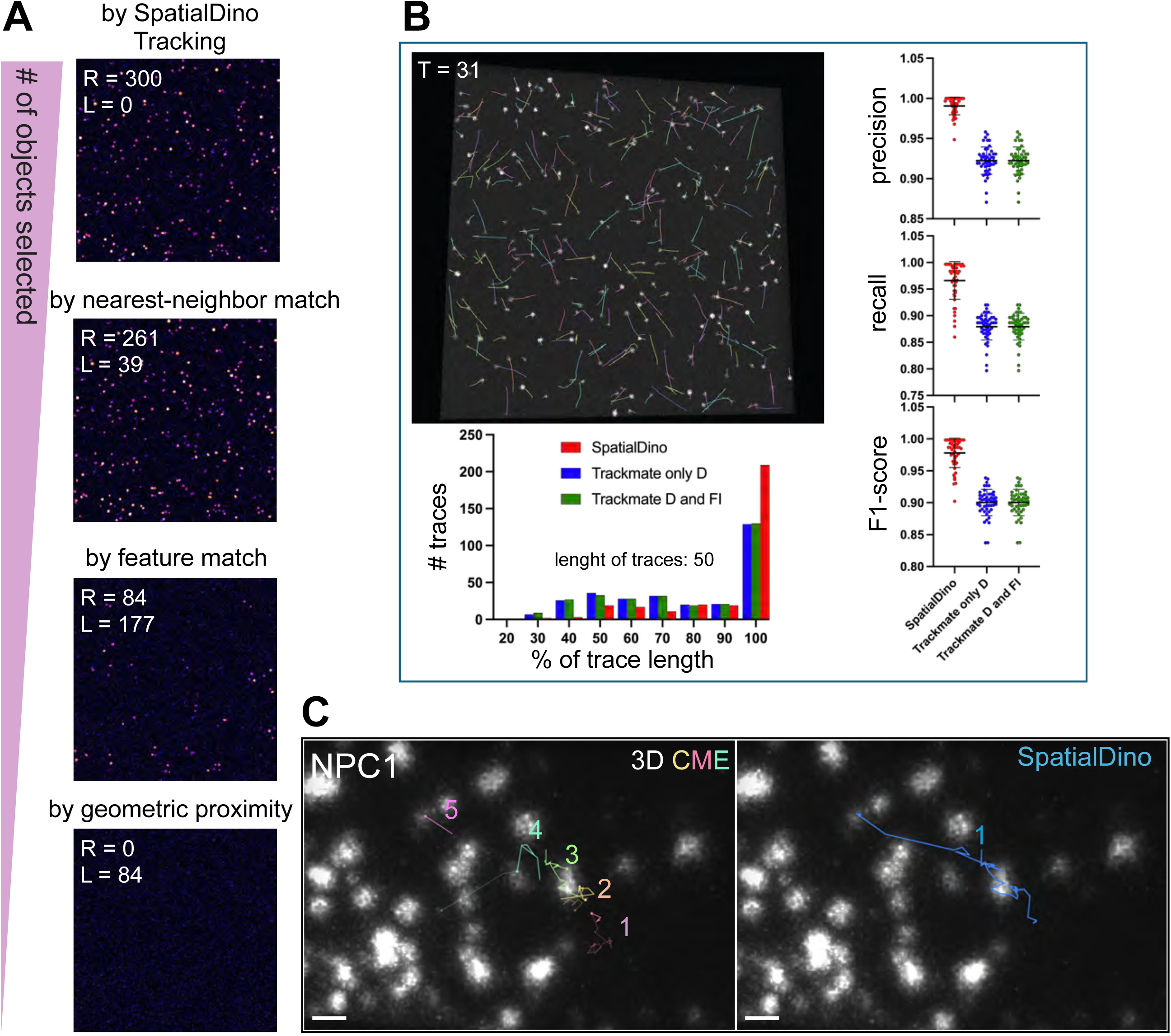
SpatialDINO enables robust 3D object tracking in time-lapse data. Associated with Videos 3 and 4. **(A)** Simulated dataset of endosomes of different sizes, shapes and intensity randomly moving in 3D. Z– maximum-intensity projections illustrate the progressive reduction in object density during sequential linking. Top to bottom: (i) initial detection of 300 objects, with remaining objects (R) and linked pairs (L) indicated; (ii) proximity-based linking within a 3-pixel radius (approximate point-spread-function width), yielding 39 linked pairs (L = 39) and 261 remaining objects (R = 261); (iii) 3D feature-similarity linking using SpatialDINO features, yielding 177 additional linked pairs (L = 177) and 84 remaining objects (R = 84); and (iv) final closest distance linking of the remaining 84 objects. **(B)** Same simulated endosome dataset used from **(A)**. Tracking performance of SpatialDINO versus TrackMate (22) on simulated endosome movies. Top, z-maximum projection of image at T=31 overlaid with trajectories reconstructed by SpatialDINO. Bottom, histogram of fractional track-length recovery, showing the distribution of trace lengths detected by SpatialDINO relative to those detected by TrackMate, scored using either distance D as the only criterion or D together with fluorescence intensity. Right, Metrics shown by dot plots include precision (fraction of links between consecutive time points that are correct), recall (fraction of ground-truth links recovered), F1 score, and the number of complete ground-truth tracks recovered. **(C)** Experimental dataset. Example comparison of tracking by SpatialDINO versus 3D CME on live-cell 3D LLSM time series (2.6-s sampling; 100 time points). Endosomes carried NPC1–Halo labeled with JFX-646, expressed by gene editing in SVG-A cells. Replicate images show a single plane at t = 0 overlaid with a cumulative, time-z–projected trajectory for one endosome: a single continuous track from SpatialDINO (left) versus four fragmented tracks from the 3D CME pipeline (21) (right). Scale bar, 1 µm.

Using the simulated endosome movies introduced in Fig. 7A, we linked detections between adjacent frames with a simple, staged procedure that progressively reduces assignment ambiguity (Fig. 8A). We first linked objects within a short-range radius consistent with the point-spread-function scale. For the remaining candidates, we linked within a larger search radius using a feature-wise voting scheme based on SpatialDINO outputs. Specifically, for each feature we identified the closest match between objects and assigned a single “win” to that pairing, independent of the magnitude of the difference. We then summed wins across features to score object–object correspondence and selected the highest-scoring link. This approach does not minimize a continuous distance in feature space and is therefore insensitive to differences in feature scale or variability that could dominate a distance-based metric. Finally, we linked any residual objects by closest geometric distance. This sequence rapidly decreased the density of unresolved candidates and preserved plausible correspondences even when objects approached closely.

To quantify performance, we compared SpatialDINO-based tracking to TrackMate (22) on simulated data comprising 300 endosomes of variable shape and low SNR randomly moving in 3D over 50 time points (Fig. 8B). Both methods detected all objects in each frame, but complete trajectories often fragmented when particles became optically unresolved during close encounters. For both approaches, most failures arose at crossing events in which objects transiently merged below the resolution limit, leading to split trajectories upon re-separation. When objects remained separable, SpatialDINO maintained identities through the feature embedding, whereas failures reflected occasional loss of identity in feature space during these unresolved intervals. Overall, SpatialDINO recovered slightly over 200 full-length tracks, whereas TrackMate recovered 135, consistent with improved link stability by SpatialDINO under crowded conditions.

We finally tested the same SpatialDINO approach on live-cell 3D LLSM time series of endosomes labeled on their limiting membrane with NPC1–Halo–JFX-646 (2.6-s sampling; 100 time points). Over adjacent frames, NPC1 content should be approximately conserved, while apparent morphology can vary with orientation and axial anisotropy. In a representative example (Fig. 8C), SpatialDINO produced a single continuous trajectory for one endosome, whereas a conventional 3D CME-based pipeline fragmented the same trajectory into five tracks of which one was incorrectly linked from another unrelated object.. Visual inspection confirmed that the SpatialDINO assignments bridged intervals that otherwise produced breaks in the tracks, indicating that feature-based identity helped resolve crowded or near-crossing events.

## DISCUSSION

SpatialDINO addresses a persistent limitation in volumetric microscopy: most analysis pipelines require task-specific training, curated ground truth, or handcrafted priors that often fail when imaging conditions or specimen types change. By adapting a self-supervised transformer strategy to native 3D fluorescence volumes, SpatialDINO produces a dense feature field that preserves the spatial organization of the underlying image while remaining agnostic to labels and downstream tasks. Our results show that SpatialDINO detects features in 3D images, particularly at medium to low SNR, better than the state-of-the-art DINOv2.

The self-supervised 3D vision transformer pipeline in SpatialDINO makes analysis of complex volumetric fluorescence microscopy images of structures across a wide range of shapes and sizes more practical and more broadly useful than currently available procedures. Though trained with a limited set of non-deconvolved single-channel fluorescence images of relatively low SNR acquired using LLSM, SpatialDINO generates features from different types of naïve 3D images of a broad type of structures including many whose characteristics and appearance were not seen during the training phase. The capacity of SpatialDINO to extract meaningful features without further training generalizes to 3D data sets acquired with other modalities of volumetric imaging, not only light sheet and spinning disc fluorescence microscopy that generate images like LLSM, but also images as different as those acquired by MRI. The ability of SpatialDINO to extract features without further training, is powerful because it reduces dependence on manual annotation while maintaining performance in complex cellular environments.

SpatialDINO extracts 390 semantic features; what this means is that each voxel that in the raw image was associated with an intensity value, is now represented by a 390-variable vector that contains semantic information relating to the intensity value of the voxel with the 3D spatial distribution of values around it. We show that the local aggregation of this contextual knowledge maps to the unique objects of different sizes and shapes present in the original image, ranging from a small spot corresponding to a point spread function to larger objects ranging from a small cluster of PSF’s (endosomes) to extended sheets (membranes) and solid volumes of variable shape (nuclei, tumors).

Knowledge of features and not just signal intensity with a given object provides a convenient and powerful tool that enables us to efficiently extract additional information for the temporal domain. Specifically, in our laboratory we are interested in tracing the position and intensity characteristics of objects as they interact and move within the environment. This has been particularly challenging to achieve, for example when tracing virus particles or endosomes and lysosomes and their cargo as they move in the relatively crowded environment inside cells. We made extensive use of 3D CME (21, 23), which builds on CMEanalysis (23) and on the pioneering u-track work of Jaqaman and Danuser (24); this approach determines valid local clusters of voxels whose intensity above background are defined by a PSF or a small cluster of them; the centroid of such objects is then subjected to tracing protocols. The approach works very well in relatively uncrowded volumes but often breaks when the trajectories of the objects collide. The features for a given object created by SpatialDINO significantly helps resolve this type of problem; tracking is now more heuristic since now the relative temporal invariance between adjacent time points in a time series in the aggregate feature values for a given object provides a very strong constraint that aids in the successful linking of the object between consecutive frames. We provide validation for this approach in two ways; by using simulated objects of different shapes and relatively low SNR signal moving in a crowded environment and using experimental data of fluorescently tagged endosomes imaged by live-cell 3D LLSM. The former allows us to strictly compare with the information embedded in the ground truth of the data; the later allows to test in a realistic experimental context. Our results demonstrate the superior performance of SpatialDINO when compared with the state-of-the-art 3D CME.

A first result is that inference alone yields a voxelwise feature volume whose principal components track object geometry and image content across disparate specimens and modalities. PCA renderings preserved puncta, crowded nuclear-envelope structures, membranes, and fine protrusions, even for image types not represented during training. This behavior suggests that the model learns generic, spatially coherent descriptors rather than memorizing classes, and that these descriptors retain information relevant to boundaries and 3D intensity distributions. PCA, however, does not by itself define object occupancy. A key advance here is that simple foreground–background likelihood mapping, computed directly from the inferred feature distributions, converts the 390-dimensional feature map into a probability volume that better represents objects in 3D. This step requires no retraining and only minimal user input—selection of representative foreground and background regions, either manually or through existing detection pipelines and can be performed on data from one dataset and applied to other, very different types of images. The probability volumes sharpen boundaries, suppress diffuse pools and deconvolution-dependent background, and generalize across image types, including naïve examples that were not used to define the foreground– background exemplars. In the tractin–GFP dataset, the probability volume recapitulated a specialized surface-reconstruction workflow, while operating directly on native 3D data and without tailoring to a particular specimen class. Together, these results position SpatialDINO as a practical front end for volumetric segmentation in settings where ground truth is unavailable or where the cost of retraining is prohibitive.

A second advance is that SpatialDINO features support instance-level identification. In simulated data representing endosomes of different shapes and signal intensity moving in 3D, object-restricted PCA signatures were effectively unique and enabled accurate instance segmentation across hundreds of objects at low SNR and high crowding. In experimental data, the same PCA-based assignment produced near-unique identities for dextran-labeled endosomes and lysosomes, and probability-map segmentation enabled reliable instance detection for Hoechst-stained nuclei. The MRI example extends the potential usefulness even further: within-volume lesion exemplars generated a probability segmentation that highlighted the contralateral lesion at the expert-annotated location, despite the absence of modality-specific training. While this single case does not replace supervised medical pipelines, it demonstrates that the feature-to-probability conversion can translate to imaging physics and texture statistics far from fluorescence microscopy.

The practical consequence of stable instance identity is improved tracking in crowded 4D data. A simple linking strategy that uses proximity first and feature similarity second increased recovery of complete tracks in simulated endosome movies compared with a widely used baseline. In live-cell LLSM time series, feature-based identity bridged intervals that fragmented trajectories in a conventional pipeline. These results emphasize that a learned, voxelwise representation can supply identity cues that remain informative when objects approach, overlap, or transiently merge below the optical resolution.

Several limitations suggest immediate directions for development of the post-processing steps following inference. Our quality assessments in many real datasets rely on visual inspection because formal ground truth is rarely available for 3D live-cell imaging; broader validation will require additional annotated benchmarks and systematic perturbations. The foreground–background probability mapping is intentionally simple; more expressive density models or spatial regularization could further improve occupancy estimates, particularly for touching objects and heterogeneous backgrounds. A limitation of the current approach is that our tracking uses a simple feature-wise voting scheme; developing more expressive linkage metrics in SpatialDINO feature space should improve robustness, especially for heterogeneous objects and variable signal. Finally, inference currently yields representations at a fixed feature dimensionality and scale; incorporating multi-scale features or explicit temporal context may improve both segmentation of complex morphologies and long-range tracking stability.

Overall, SpatialDINO provides a general-purpose, self-supervised route from raw 3D images to dense, spatially faithful embeddings that can be converted, using minimal post-processing into probability volumes, instance identities, and robust trajectories. The progress is practical as well as conceptual: a single model supports segmentation and tracking across multiple microscopy modalities and even extends to MRI, reducing dependence on task-specific retraining while preserving the fine structure needed for quantitative cell biology.

## MATERIALS AND METHODS

### Training dataset

We trained on single-channel fluorescence time series acquired by 3D lattice light-sheet microscopy (LLSM) from living cells. The time series were a sampling of data acquired in our laboratory and were not collected specifically for the purpose of creating the training dataset. Each time point comprised a volumetric image with variable morphological and structural complexity. The training dataset comprised 78 independent imaging experiments, each generating a live-cell 3D LLSM time series. In total, the raw dataset contained 45,461 volumetric stacks, ∼2.4 TB after curation.

The dataset consisted of gene-edited human cells expressing fluorescent markers for endocytic structures. SVG-A cells expressed the early endosomal marker EEA1–mScarlet and the late endosomal/lysosomal marker NPC1–Halo, labeled with JFX-AF64 or JF646; it also included imaging data from SUM159 cells and SVG-A cells incubated with fluorescent dextran (AF-647) to label late endosomes and lysosomes by uptake and accumulation. The dataset also included gene-edited SUM159 cells expressing Ubash-3B–Halo labeled with JF646. These structures ranged from single diffraction-limited puncta to small clusters. Within a given 3D volume, many puncta were stationary, but between successive time points they exhibited variable 3D displacements. To expand the range of morphologies, we imaged EEA1-and NPC1-expressing cells after treatment with the PIKfyve inhibitor apilimod, which enlarges NPC1-positive late endosomes (11) . Under these conditions, NPC1 labeling outlined an expanded limiting membrane rather than filling a compact object, providing examples with membrane-defined boundaries.

Together, these 3D time series spanned broad ranges of fluorescence intensity, signal-to-noise ratio, object density, and shape heterogeneity, from relatively isolated puncta to more crowded volumes with partially overlapping endosomes, reflecting the diversity encountered in routine live-cell imaging.

#### Data curation

A typical single-channel 3D LLSM volume contained ∼10^8 voxels (1000 × 1000 pixels × 100 planes; 16-bit) and required ∼2 × 10^8 bytes (∼200 MB) of storage. To collect tens of thousands of training examples, we acquired data as live-cell movies. Each movie comprised a time series (T + XYZ) of typically 50 time points with two or three fluorescence channels captured a broad range of morphologies and subcellular structures.

During early development of the SpatialDINO training workflow, we found that efficient training required two steps: suppressing spurious noise and restricting learning to regions that contained relevant cellular signal. Manual curation was infeasible at this scale. We therefore implemented the semi-automated filtering and validation pipeline described below to retain biologically informative volumes while reducing overall dataset size.

#### Semi-automated filtering and validation pipeline

For each channel in each 4D training movie, we computed maximum-intensity z-projections and used them for rapid visual screening. This step was used to reveal acquisition defects and allowed us to exclude corrupted or low-quality volumes before downstream processing (Fig. 3).

#### K-means cropping

A large fraction of our intracellular 3D volumes contained mostly background. Consequently, the random cropping commonly used for natural images often yielded crops dominated by background when applied to sparse microscopy data. We therefore implemented *k-means cropping*, a sampling strategy that enriched for data-dense regions within each volume. We used k-means clustering to identify k regions of high signal density (empirically, k = 4 for our data). The pipeline automatically extracted and saved region-of-interest crops on the corresponding volumes deskewed to correct the geometric displacement introduced by oblique LLSM acquisition; these validated volumes served as the training data.

Because voxel intensities spanned a wide dynamic range, we first applied percentile-based thresholding followed by median filtering to generate a robust binary mask that remained relatively insensitive to outliers and imaging artifacts. We sampled points from this mask and fit k-means; we used the resulting centroids as crop centers. We then sampled crops around each centroid, with side lengths set to 0.3–0.6 of the corresponding volume dimensions (chosen empirically to capture complete structures while limiting background). Relative to random cropping, k-means cropping yielded crops containing ∼3–5× more biological signal and improved training efficiency.

Beyond signal enrichment, k-means cropping improved the learned feature representations (see SpatialDINO, [specify section]). Although we developed this procedure for fluorescence microscopy, the same strategy could be applied to other volumetric imaging modalities, including natural-scene volumes, when informative content occupies a small fraction of the field of view.

#### K-means cropping pseudocode

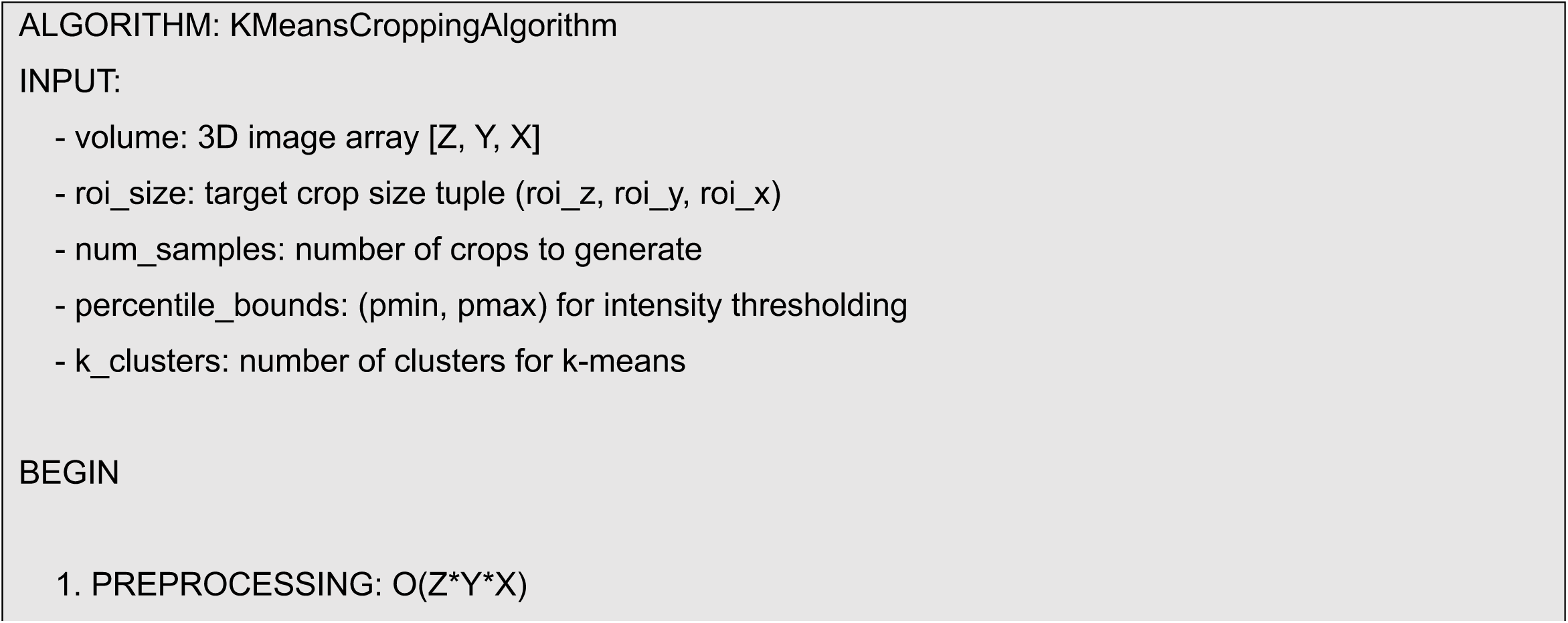
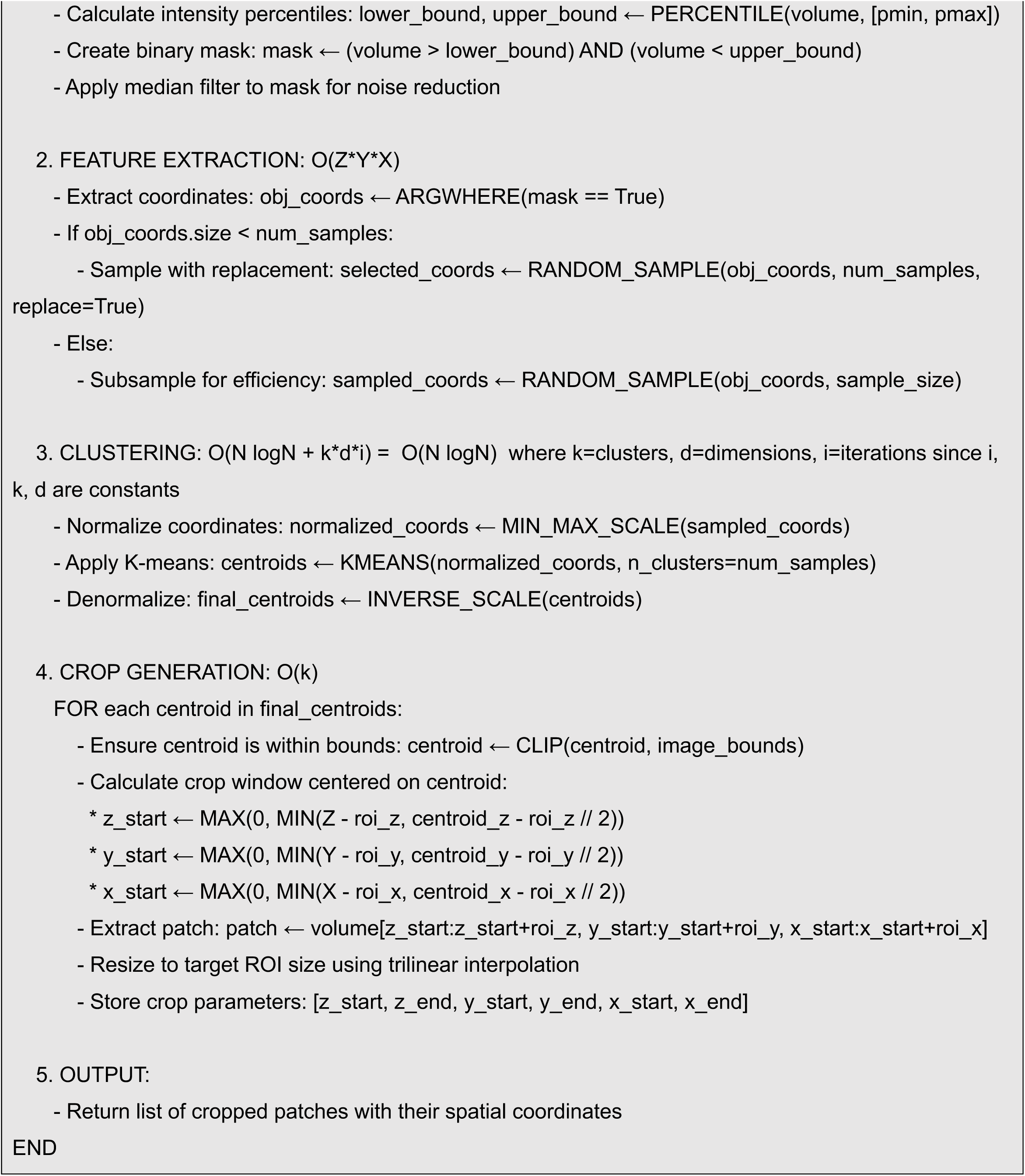

#### Compression of the dataset for training

The curated dataset comprised many small files containing cropped ROIs and associated metadata, which imposed substantial I/O overhead during training. To reduce I/O load, we stored the samples in WebDataset format (25) and packaged them into POSIX tar archives (“shards”) of ∼950 MB each. Each shard contained ∼60 single-volume, single-channel samples (see File System Optimization). Using distributed processing across a three-node cluster (256 CPU cores per node), we converted ∼2.4 TB of curated data into ∼2,600 shards in ∼12–14 h. We provide this dataset publicly (see Code and Data Availability).

### Inference dataset

For inference, we used datasets that were not used for training and that span multiple imaging modalities. The fluorescence microscopy datasets include cells expressing AP2 as a fluorescent marker for endocytic clathrin-coated pits and vesicles; NPC1, a constitutive marker of late endosomes and lysosomes; and internalized fluorescent dextran to label late endosomes and lysosomes. We also included Hoechst-labeled nuclei and cells expressing plasma-membrane markers to visualize cell-surface structures. To test cross-modality transfer, we further analyzed a breast MRI volume from a human subject with a malignant lesion. Experimental details for each dataset (specimen, labeling, imaging modality, and acquisition parameters when available) are given in the corresponding Results sections and in the relevant figure and movie legends.

### Model architecture

#### DINOv2 overview

Self-supervised learning (SSL) aims to learn representations from unlabeled data that support downstream tasks (26). Without labels, SSL uses indirect objectives (“pretext tasks”) derived from the data. These objectives typically bring representations of augmented views of the same image (e.g., masked, noised, or rotated) closer together while separating representations from different images. Joint-embedding methods enforce this objective directly in representation (latent) space, whereas reconstruction-based methods optimize in pixel space through a decoder. Reconstruction objectives often train more stably, but joint-embedding approaches can yield more informative features, particularly under noisy conditions (27, 28), and thus suit everyday fluorescence microscopy datasets.

A central failure mode of joint-embedding SSL is representation collapse, in which the model maps all inputs to the same representation. Preventing collapse requires a mechanism that maintains separation among representations from different samples. Explicit contrastive strategies use negative samples but generally demand very large batch sizes, which can be impractical for large datasets. Alternatively, methods can discourage collapse implicitly through carefully chosen augmentations, regularization, and hyperparameter tuning (29).

SpatialDINO model is based on the DINOv2 family (30), which provides state-of-the-art joint-embedding SSL for images (see Fig. 1A for a schematic representation of the framework). DINOv2 processes global and local views of each input (large and small augmented crops) using a teacher–student pair. The student is trained to match the teacher outputs in representation space. The teacher is updated by self-distillation, using an exponential moving average (EMA) of the student parameters from preceding iterations. To avoid collapse (31), DINOv2 combines several stabilizing components, including projection heads (weight-normalized linear layers), centering of the teacher softmax outputs, Sinkhorn–Knopp normalization, and KoLeo regularization, with tuned hyperparameters (29, 30). This design yields both global and spatially dense representations suitable for downstream tasks.

Architecturally, DINOv2 uses a Transformer encoder (32), following the Vision Transformer (ViT) formulation (33). The input RGB image is partitioned into nonoverlapping 14 × 14 patches; each patch is flattened and mapped to a higher-dimensional *patch token*. Positional encodings are added to retain 2D spatial information. The token sequence is augmented with a learned single classification [CLS] token to aggregate global image information and with register tokens, introduced subsequently in the DINOv2 codebase, to reduce the compressive burden on the [CLS] token and thereby mitigate high-norm artifacts in feature maps (34). Patch tokens and the [CLS] token feed the DINO (35) and iBOT (36) projection heads, respectively. At inference, we discard the projection heads and only use the transformer encoder outputs.

DINOv2 optimizes three loss terms: a DINO loss (cross-entropy between student and teacher outputs from the DINO heads) that drives global, image-level features; an iBOT loss of analogous form that guides dense, patch-level features; and a KoLeo term that provides regularization critical for preventing collapse (29, 30).

#### SpatialDINO: extending DINOv2 for 3D spatial representations

Most adaptations of DINOv2 to volumetric data use a “2.5D” strategy: they apply a 2D model independently to each z plane and then aggregate the outputs into a 3D feature volume (37). This approach conveniently reuses pretrained 2D weights, but it can introduce plane-to-plane discontinuities and limits expressivity by discarding true volumetric context. We therefore adapted DINOv2 to operate natively in 3D, replacing plane-wise processing with a volumetric tokenization and training scheme that preserves cross-plane context. The resulting architectural and training modifications define SpatialDINO (Figure 1 schematizes the SpatialDINO pipeline).

To move from 2D to 3D while preserving the core ViT design, we replaced the 2D patch-embedding layer with a 3D convolution-based patch embedder that converts input volumes into token sequences compatible with the ViT backbone. We also adapted data augmentation to 3D microscopy. We replaced random global/local cropping with k-means cropping to avoid background-only views. Photometric transforms designed for natural RGB images performed poorly on grayscale single channel fluorescence microscopy intensities and were omitted. We instead applied random contrast adjustment, random 90° rotations, and additive Gaussian noise (global views only), each of which yielded modest performance gains.

Fluorescence microscopy places greater weight on high-frequency spatial information than do many natural-image tasks. The original DINOv2 patch size (14 voxels) produced overly smooth features for small structures, degrading segmentation of diffraction-limited objects such as clathrin coated pits or vesicles and viruses. We therefore used an 8-voxel patch size, balancing spatial resolution against computational cost.

Training a 3D ViT using the DINOv2 strategy without collapse would have required batch sizes beyond our available compute. In practice, training runs often collapsed after ∼10,000 iterations, producing high-norm tokens and grid-like artifacts that degraded feature maps. We attribute this susceptibility to two factors: (i) a single [CLS] token compresses substantially more information in 3D, and (ii) compared with natural images, our volumes show reduced inter-sample diversity (diffraction-limited structures, grayscale intensities, and noise), which narrows the embedding distribution and promotes collapse. Register tokens reduced, but did not eliminate, these failures. We prevented collapse by removing positional encodings (NoPE) (18, 19).Remarkably, even without explicit positional encodings, the attention layers learned useful positional structure, consistent with earlier reports in 2D ViTs (20). We also observed a small improvement in feature-map quality by adding a single zero-initialized register token at inference, as proposed in (38).

Finally, to reduce memory footprint, we decreased the output dimensionality (“number of prototypes”) of the DINO and iBOT heads from 131k to 32k without measurable loss of representation quality. Our final model used a ViT-small backbone with ∼21M trainable parameters.

### Training Pipeline

We trained on 3 DGX-A computers each with 8 NVIDIA A100 GPUs (40 GB; see Hardware Specifications) for 250,000 iterations (∼4 d) using the configuration parameters listed in Supplementary Table 1.

#### Data Pre-processing

During pre-training, after loading volumetric crops from the extraction pipeline, we corrected anisotropic voxel spacing by trilinear interpolation with scale factors 1.0, 1.0, 2.404) in X, Y and Z, respectively, as determined by our LLSM imaging geometry. We then applied k-means cropping again to generate data-centric sub volumes. From these, we generated local and global views and applied corruption transforms independently to each view for the pretext tasks, including random flips along all three axes, random rotations, random contrast adjustment, and additive Gaussian noise (global views only). We used a batch size of 14. Each sample comprised two global crops (112 × 112 × 112 voxels) and eight local crops (48 × 48 × 48 voxels).

#### File System Optimization

Training 3D vision transformers on volumetric data stressed the storage stack. Our training cluster comprised three NVIDIA A100 compute nodes connected by InfiniBand (10 GB/s). The dataset spanned terabytes and comprised many small samples; under these conditions, conventional network file systems (NFS) could not sustain the concurrent, low-latency reads required to maximize GPUs throughput.

In initial runs with standard NFS, concurrent access from many GPU workers produced severe I/O bottlenecks, with batch read latencies >500 ms. NFS imposed a single-server bottleneck: all workers shared one network interface, leading to request queuing and cache thrashing. We improved NFS performance by enabling Remote Directory Memory Access (RDMA) over the InfiniBand protocol rather than TCP/IP as the default transport, which reduced sequential read latency by ∼2×. While these changes improved throughput, they did not remove the architectural limitation of NFS under high-concurrency, small-file workloads. We therefore stored the preprocessed crops in WebDataset format. Packaging samples into POSIX tar archives (“shards”) enabled streaming reads without per-sample file opens and metadata lookups. This shifted access toward large sequential reads, which NFS serves efficiently, and yielded sustained aggregate read rates of ∼1.8 GB/s across the cluster (∼60% improvement relative to per-file access). Shards also partitioned naturally across workers and nodes: each worker claimed exclusive shards, reducing contention and ensuring full dataset coverage during distributed training. Compatibility with standard tar utilities simplified inspection and debugging.

#### Computational Optimizations

Training a 3D vision transformer on volumetric fluorescence microscopy data required optimization across the software and hardware stack. Transformer self-attention scales quadratically with sequence length; for our 3D token sequences, this increased compute substantially relative to comparable 2D inputs (approximately eightfold for typical volume sizes).

We used mixed-precision training with BF16 (39) to reduce activation and parameter memory by ∼50% while retaining numerical stability relative to FP16 because of BF16’s wider exponent range. We nevertheless computed the DINO and iBOT heads in FP32 during pre-training to preserve precision in the projection and loss computations. For distributed training, we used PyTorch Distributed Data Parallel (DDP) (40) rather than Fully Sharded Data Parallel (FSDP) (41). Given our model size and hardware, DDP provided the best trade-off between implementation complexity and performance, whereas FSDP offered only marginal gains.

#### Scaling Implications

The architectural decisions and optimizations implemented for SpatialDINO training have significant implications for scaling larger models and datasets in biomedical imaging. Our current infrastructure successfully trained a 21M parameter ViT-Small model (30), but the biomedical community’s trajectory toward foundation models with billions of parameters necessitates careful consideration of scaling strategies. The quadratic scaling of scaling *O*(*n*^2^) transformer memory requirements with sequence length becomes particularly challenging for 3D data. This suggests that future scaling will require either architectural innovations like hierarchical transformers sub-quadratic attention usage or hardware advances such as multi-GPU model parallelism.

The success of our WebDataset-based I/O optimization demonstrates that resourceful data formatting can overcome traditional bottlenecks, but scaling to petabyte-sized datasets will require true distributed file systems. Our projections indicate that training on 50× larger datasets (approaching 100TB) would saturate even our optimized NFS setup, necessitating migration to systems designed for AI workloads.

Looking toward the future of biomedical foundation models, our experience with SpatialDINO provides a roadmap for infrastructure requirements. The most important scaling insight from our work is that domain-specific optimizations, from our fluorescence microscopy-adapted preprocessing to our dimensionality reduction in tracking provide multiplicative improvements that can offset the need for pure computational scaling. This suggests that the path to powerful biomedical AI models lies not just in larger clusters, but in imaginative co-design of algorithms, systems, and domain knowledge.

### Ablation Studies

We implemented SpatialDINO by systematic ablation of data processing, model architecture, and loss configuration. Unless noted otherwise, we pre-trained each variant for 1,000 steps using a fixed random seed (0) and compared it with the baseline configuration. For the crop-extraction method ablation, we trained for 200,000 steps to capture longer-term effects (see Crop Extraction below). When a configuration exceeded GPU memory, we used gradient accumulation to match the baseline effective batch size.

Because no benchmark 3D dataset exists for our specific application, we evaluated representation quality qualitatively. For each ablation, we inspected one or both of the following readouts obtained using a naïve endosome image that had not been used during training: (i) principal component analysis (PCA) of the patch-token features and (ii) the summed attention weights between the [CLS] token and patch tokens across all heads in the final attention layer (Fig. S1). We interpret patch tokens as encoding feature-specific content and the [CLS]→patch attention map as a compact descriptor of spatial structure (often resembling a coarse pseudo-segmentation). For visualization, we rendered min- and max-intensity z projections of the resulting feature volumes. For these analyses, we input the encoder with the original-resolution volume up sampled 2×, without isotropic rescaling and without chunked attention.

#### Intensity Normalization

Fluorescence microscopy 3D images require intensity normalization strategies that differ from those used for natural images. We evaluated several approaches and ultimately adopted a histogram-based normalization followed by min–max scaling.

We first applied simple min–max normalization. This approach proved inadequate in the presence of microscopy-specific artifacts (e.g., hot pixels, dead pixels, sporadic local signal saturation), which produced spurious extrema and compressed the dynamic range of the remaining signal.

We next tested min–max normalization followed by ImageNet mean–variance normalization (42). This procedure introduced an inductive bias that improved segmentation performance, consistent with partial transferability of natural-image statistics to microscopy data. However, ImageNet normalization required expanding the single-channel volume to three channels, imposing an artificial color structure and tripling data size. Moreover, it did not capture the intensity distributions of fluorescent puncta in a principled way. We therefore did not adopt this strategy, but we note it as a potentially useful direction for future work.

Third, we applied min–max normalization followed by z-score standardization using the mean and SD estimated from our training set, analogous to ImageNet-style preprocessing (42). This approach also performed poorly. It relied on stable global statistics across a heterogeneous dataset and often attenuated faint particles with intensities below the dataset mean.

Finally, we chose a histogram-based normalization followed by min–max scaling to [0, 1]. This procedure preserved usable dynamic range while remaining insensitive to rare outliers and to the strongly skewed intensity distributions typical of fluorescence microscopy. This method relates conceptually to histogram-based contrast methods such as contrast-limited adaptive histogram equalization (CLAHE) (43), but we implemented it as a global, volume-wise normalization. We binned 16-bit intensities into 65,536 histogram bins and defined lower and upper bounds as the first and last intensity values for which the bin count exceeded (1/5000) × (total voxels). We then clipped intensities to these bounds and rescaled to [0, 1].

#### Crop Extraction

Compared to baseline random cropping, our k-means cropping during pre-training improved the quality of model features, by shifting the attention value distribution to higher values for foreground and lower values for background. We attribute this to the fact that the views of data it produced had more foreground prominence, for which the model could reserve more capacity instead of on unimportant background information.

Interestingly the improvements of k-means cropping compared to our baseline could not be seen in the attention features until the 200000-step mark in training where the model shifted from memorization to generalization. We attribute the cropping strategy to be a significant factor in the development of emergent behavior in self-supervised learning methods such as DINOv2 but under-discussed in literature. ImageNet-based datasets with dominant foreground objects lead to supreme results compared to other datasets because of the models learning signal being adversely affected with less informative crops of foreground with basic cropping methods. This sampling strategy proved essential for efficient self-supervised learning in the sparse volumetric domain where random cropping would lead to over-representation of background as well as translates to diverse dataset types.

#### Crop Sizes

We evaluated the impact of crop dimensions on model performance and computational efficiency. Larger local and global crops demonstrated improved performance by providing more spatial context for learning volumetric representations but came at significant computational cost. The memory requirements scaled cubically with crop size, making the largest crops impractical for multi-GPU training with our available hardware.

Initially, our Z dimension of crops was half of the Y and X to account for the discrepancy in data density, however, we identified that this limited the patches of Z context, which hurt the representation quality. As a result, we investigated cubic crop sizes, so the model would be able to learn effective representations.

Notably, we adopted roughly the same ratio for global and local crop dimensions (∼ 2.33) as used in DINOv2. We also kept the same number of global and local crops, two and eight respectively.

We compared the performance of the following configurations: 112 x 112 x 112 for global crop size and 56 x 56 x 56 for local crop size, versus 224 x 224 x 224 for global crop size and 98 x 98 x 98 for local crop size. We found a balance between computational cost and performance with a global crop size of 112 x 112 x 112 and local crop size of 56 x 56 x 56 effectively providing sufficient contextual information and capturing fine-grained features.

#### Masking

We compared the masked autoencoder (MAE) random masking strategy (44) with iBOT’s block masking (36) to identify an effective masking scheme for 3D volumes. Independent random voxel masking provided a weak learning signal in our data, because neighboring voxels in 3D fluorescence microscopy data are highly correlated. Under this regime, the network could often infer masked voxels by local interpolation, reducing the difficulty of the pretext task.

Block masking, which removes contiguous 3D regions, better exploited spatial dependence to create a harder reconstruction problem. It forced the model to use larger spatial context and to learn structural regularities, yielding more robust representations. Block masking also allowed us to bias masks toward foreground, so that most masked voxels contained biological signal rather than background. This increased the density of informative masked targets during self-distillation. We implemented iBOT-style masking by extending its 2D block masking to randomly positioned 3D blocks and used the same relative block-size proportions as in the 2D formulation (36).

#### Stack versus volume patch processing

We evaluated two approaches to 3D fluorescence microscopy inputs: DINO2.5D REF, which processes each z slice (or z-stack plane) independently and then aggregates representations, and SpatialDINO, which tokenizes and processes the volume directly as 3D patches. The 2.5D approach discards inter-plane dependencies and can lose 3D structural context. In contrast, native 3D patch processing preserves volumetric relationships that matter for cellular morphology. In our experiments, SpatialDINO consistently outperformed the stack-based approach, with the largest gains for structures spanning multiple z planes.

#### Patch size

We compared patch sizes of 8 voxels and 14 voxels (the DINOv2 default) to balance spatial resolution against computational cost. Because many biologically relevant objects in our data are small, we expected smaller patches to better preserve the features required for accurate segmentation. With a patch size of 14, the encoder produced representations that were too coarse to resolve small structures such as clathrin coated pits or coated vesicles and viruses. Reducing the patch size to 8 substantially improved resolution of these objects while keeping compute tractable. Smaller patches improved performance further but at prohibitive cost. We therefore selected an 8-voxel patch size as a practical Pareto-optimal trade-off between feature resolution and efficiency.

#### Positional encodings

We compared three positional encoding strategies: learned absolute embeddings, fixed sinusoidal absolute encodings, and no positional encoding (NoPE) (19). In our 3D setting, positional encodings consistently produced grid-like, high-norm artifacts in feature maps and were associated with collapse after extended training (>10,000 iterations), often accompanied by high-norm patch tokens and degraded spatial features. By contrast, removing positional encodings eliminated these artifacts and improved stability. Despite the absence of an explicit positional encoding layer, the model learned relative and absolute positional structure through attention, exploiting the inherent spatial organization of the input. Thus, NoPE reduced positional bias in long token sequences and prevented collapse while preserving spatial organization in the learned features. This behavior parallels recent observations in 2D vision transformers (20) and is particularly important for 3D volumes, where artificial grid patterns directly impair segmentation quality. To our knowledge, this work provides the first formal ablation of positional encodings in 3D vision transformers.

#### Register tokens

In early training runs, models frequently collapsed after ∼10,000 iterations. This instability reflected a combination of auxiliary-head loss dynamics and the grid-like artifacts associated with positional encodings. We also hypothesized that a single [CLS] token incompletely captured 3D volumetric content. In contrast to ImageNet, where natural images tend to occupy well-separated regions of latent space, our microscopy volumes are often visually similar, with blurred boundaries and recurrent high-intensity regions that can concentrate embeddings and promote collapse. We therefore tested adding register tokens during training (34). Increasing the number of register tokens produced a modest improvement in stability but did not prevent collapse. Because register tokens were originally introduced to reduce high-norm artifacts, an effect we traced largely to positional encodings, we ultimately shifted from training-time register tokens to test-time register tokens.

We next evaluated zero-initialized register tokens at inference, as proposed in (38), testing 0, 1, 2, 4, and 8 tokens. Adding a single zero-initialized register token improved segmentation quality, increasing particle detection and sharpening boundary delineation, particularly in densely packed regions that require precise instance separation. Increasing the number of test-time register tokens beyond one yielded little additional benefit while increasing compute. Because test-time register tokens require no retraining and add only incremental attention computation, we adopted one zero-initialized register token at inference as an efficient and reproducible improvement. These results suggest that a single additional token provides sufficient test-time “memory” capacity for our volumes and motivates further exploration of token-based mechanisms for robustness.

#### Prototype dimensionality reduction

To reduce memory use under 3D workloads, we evaluated smaller prototype sets for the DINO and iBOT heads. DINOv2 uses K = 131,072 prototypes for 2D images; we tested *K* ∈ {32768, 65536, 131072} for volumes. Prototype assignment probabilities follow *p* = *softmax*(*z^T^ W* / *τ*), where z is the feature vector, *W* ∈ ℝ^*D* × *K*^ is the prototype matrix, and τ is the student temperature. Reducing K from 131,072 to 32,768 produced no observable loss in representation quality by our qualitative readouts. This fourfold reduction decreased memory consumption by ∼12 GB and improved GPU utilization without compromising performance.

#### Complementary Loss Functions

We ablated the DINO and iBOT objectives to assess their individual contributions to 3D representation quality. We optimized the combined objective

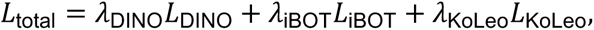

with the KoLeo regularizer

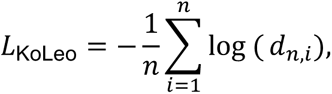

which promotes feature diversity. We evaluated three settings: (i) *λ*_DINO_ = 1, *λ*_iBOT_ = 1; (ii) *λ*_DINO_ = 0, *λ*_iBOT_ = 1; and (iii) *λ*_DINO_ = 1, *λ*_iBOT_ = 0. Using iBOT alone reduced feature quality; using DINO alone produced a similar reduction. Thus, the two terms act complementarily. iBOT provides patch-level supervision through masked-token prediction, whereas DINO enforces consistency of the global [CLS] representation across augmented views (*p_t_* ≈ *p_s_*), stabilizing the volume-level embedding.

#### Pixel versus latent reconstruction

We tested whether adding pixel-level reconstruction improves representations by adapting a masked autoencoder (MAE) to 3D. We appended a decoder head to the encoder and reconstructed masked patches.

Given an input volume *V* ∈ ℝ^*C*×*Z*×*X*^ partitioned into nonoverlapping patches, we masked a subset M comprising 75% of patches, as in (44). We evaluated both unnormalized and normalized reconstruction losses. For the normalized case,

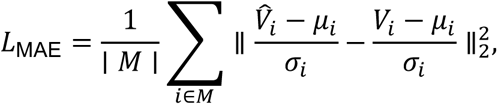

where 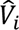 is the reconstructed patch, *V_i_* is the original patch, and *μ_i_*, *σ_i_* are patch-wise mean and standard deviation.

Although MAE-style reconstruction adds a self-supervised signal, the pixel decoder tended to reproduce noise present in the input, and the resulting features were less useful by our qualitative readouts. We therefore used latent-space objectives only, implemented through DINO and iBOT.

### Inference Pipeline

SpatialDINO inference (Fig. 2) operates on volumetric datasets that exceed the scale assumed by standard ViT implementations. A typical LLSM 4D movie contains volumes of ∼100–200 × 1024 × 1024 voxels. With an 8-voxel patch, one volume produces ∼4.1 × 10^5 tokens; after upsampling to support high-resolution segmentation, token counts rise into the millions, making full-volume inference impractical. We therefore built the inference pipeline around three components: (i) GPU-resident, chunked interpolation; (ii) multi-node execution to match the throughput of typical experiments; and (iii) a custom chunked attention module.

#### Chunked interpolation

Preprocessing, not encoding, initially limited throughput. Interpolation is required for two reasons: to resample anisotropic LLSM volumes to isotropic spacing for feature extraction, and to upsample volumes to obtain higher-resolution feature maps for segmentation. Naïve trilinear interpolation of a 200 × 1024 × 1024 volume to isotropic spacing can require >150 GB of GPU memory for intermediate tensors. We therefore implemented chunked trilinear interpolation that is mathematically equivalent to full-volume interpolation but evaluates it on small windows to bound peak memory. We exploit separability and process the volume in (k, k, k) windows, limiting intermediate tensor sizes and enabling isotropy correction and upsampling within available GPU memory.

#### Multi-node execution

The implementation supports multi-node inference, allowing concurrent processing of volumes and improved utilization of available GPUs. This design scales directly with GPU count and supports long 4D movies.

Chunked interpolation. Preprocessing, not encoding, initially limited throughput. Interpolation is required for two reasons: to resample anisotropic LLSM volumes to isotropic spacing for feature extraction, and to upsample volumes to obtain higher-resolution feature maps for segmentation. Naïve trilinear interpolation of a 200 × 1024 × 1024 volume to isotropic spacing can require >150 GB of GPU memory for intermediate tensors. We therefore implemented chunked trilinear interpolation that is mathematically equivalent to full-volume interpolation but evaluates it on small windows to bound peak memory. We exploit separability and process the volume in (k, k, k) windows, limiting intermediate tensor sizes and enabling isotropy correction and upsampling within available GPU memory.

#### Chunked attention

During inference, our typical volumes produced token sequences that exceeded GPU memory. A common workaround is to split the volume into smaller, overlapping subvolumes, run each subvolume independently, and then stitch the outputs. This sliding-window strategy works well for CNNs but fails for SpatialDINO because attention uses softmax normalization: each subvolume normalizes its attention weights independently, so adjacent regions acquire systematically different feature statistics regardless of overlap. The resulting feature volume appears as a patchwork of incompatible inference tiles.

We avoided partitioning the input volume and instead chunked attention inside the transformer, at the token level. In a standard transformer, the patch embedding converts the input into a 1D token sequence, which then enters the transformer blocks. The dominant memory and compute cost arises in self-attention (Fig. 2B).

For each head, the model projects the tokens to queries (Q), keys (K), and values (V); forms the score matrix *QK*^T^, whose size scales quadratically with sequence length; applies a softmax; and multiplies the result by *V* to produce the attention output, which then passes to the feed-forward layers. In this implementation (Fig. 2C), we partitioned the Q, K, and V sequences into non-overlapping chunks of configurable size. We then computed attention by iterating over query chunks and, for each, scanning all key/value chunks. We evaluated the softmax online with a numerically stable running-maximum scheme: for each head and query position, we maintained a running maximum *m*, a running normalizer *l*, and an accumulated numerator out in fp32. For each KV chunk, we updated *m* with the chunk-wise score maximum, rescale the existing accumulator by exp (*m*_prev_ − *m*_new_), and accumulate exp (scores − *m*_new_) into both *l* and out. After the KV scan, the attention output equals out/*l*, matching full softmax attention exactly, while bounding memory by the chunk sizes rather than the full token sequence length.

In our implementation, we precompute keys and values once per block and store them separately on CPU, GPU, or disk, depending on user constraints thereby enabling long-context inference with bounded VRAM. We implemented the online softmax update as a fused Triton kernel that tiles over heads and query blocks and performed score computation, max reduction, exponentiation, normalizer updates, and value accumulation in a single GPU launch. This design preserves full-context attention while minimizing intermediate storage and maintaining high throughput. We also evaluated the feed-forward networks (FFNs) in chunks, which bounds the transformer’s intermediate memory by the configurable chunk size rather than the full sequence length.

We used chunked attention only at inference. It does not alter SpatialDINO training or outputs and remains fully compatible without retraining. When standard inference fits in GPU memory, the conventional full-attention path remains available.

### Downstream Tasks

This section describes the downstream analyses applied to the probability maps, including object segmentation and, where applicable, tracing of spatio-temporal dynamics. We segment objects of interest by assigning each voxel a foreground confidence score that aggregates evidence across multiple 3D feature volumes. For each feature, we compare the voxel intensity to empirically learned foreground and background intensity distributions derived from an AP2-based reference segmentation (or the corresponding reference mask in other modalities) and convert these likelihoods into a per-feature foreground posterior. We then sum per-feature posteriors to obtain a voxel-wise confidence map that preferentially highlights voxels whose feature signatures match the reference foreground statistics and suppresses background-like voxels. We use this continuous-valued map as the common input for subsequent steps, including object delineation and, when relevant, trajectory construction over time.

#### Segmentation using PCA

Three-dimensional PCA volumes obtained from SpatialDINO (used for simulated and experimental endosomes) were background-subtracted with a spherical structuring element (radius, 10 voxels). We filtered the resulting images with a three-dimensional Laplacian-of-Gaussian (LoG) kernel (3 × 3 × 2 voxels), inverted intensities, and applied Voronoi–Otsu labeling to segment objects.

#### Probability map construction from empirical foreground/background intensity models

SpatialDINO acquired semantic segmentation capability through self-supervised training, without voxel-level annotations. We performed zero-shot segmentation on naïve volume images using the encoder’s dense feature maps. Each spatial patch in the input volume maps to a 390-dimensional feature vector that captures local morphology and texture.

In practice, these features increase the effective signal-to-noise ratio of the input relative to raw intensities, enabling segmentation procedures to succeed even when intensity-based segmentation failed. Empirical simple visual evaluation showed that clustering in feature space separated not only the biological classes included in the training set and membrane outlines of different shapes such as single and small clusters of diffraction limited objects, but also those not present in the training set, such as elongated structures (mitochondria) and relatively large solid objects (nuclei and tumors). This behavior indicates that self-supervised training on diverse microscopy volumes yields representations that align with biologically meaningful categories.

*1. Voxel-wise class definition.* We defined voxel-wise class membership using a binary 3D segmentation mask. The mask assigned each voxel to either **foreground** (object-associated voxels) or **background** (all remaining voxels). For experimental cell datasets, we segmented clathrin-coated pits using the AP2 reference channel (fluorescently tagged AP2) and a local-maxima–based detection procedure using 3D CME (21), yielding a binary foreground/background partition. We pooled all foreground voxels into a single class for probabilistic modeling and did not model individual objects separately. To ensure consistent feature-based inference across acquisitions, we applied the same binary class definition framework to other imaging datasets (e.g., raw lattice light-sheet, spinning disk, deconvolved, and simulated volumes), using masks aligned to each dataset’s voxel grid. For the breast cancer MRI example, we defined a tumor foreground mask by thresholding intensity and assigned all non-tumor voxels to background.

*2. Feature volume generation and alignment.*

For each dataset, we generated *N*_feat_ = 390 three-dimensional feature volumes, each representing a distinct voxel-wise intensity map derived from the raw imaging data. We maintained voxel-to-voxel correspondence between each feature volume and the segmentation mask by aligning them on the same (*x*′ *y*′ *z*) grid without rescaling or interpolation, thereby preserving spatial fidelity.

*3. Empirical class-conditional intensity distributions per feature*

For each feature *i*, we estimated empirical intensity distributions for the two classes by sampling intensities from all voxels assigned to background and foreground by the segmentation:

- 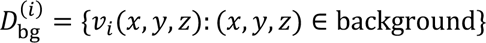
- 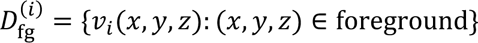

We imposed no parametric assumptions (e.g., normality or unimodality) on these distributions. Instead, we fit kernel density estimators (KDEs) to obtain continuous class-conditional density functions:

- 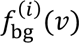 for background
- 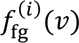 for foreground

*4. Per-feature voxel-wise posterior probabilities.*

For each voxel at (*x*′ *y*′ *z*) and each feature *i*, we evaluated the KDEs at the observed voxel intensity *v_i_*(*x*, *y*, *z*) to obtain class likelihoods:

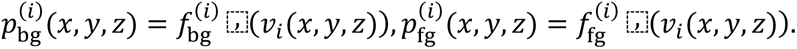

We converted these likelihoods into two-class posterior probabilities by normalization:

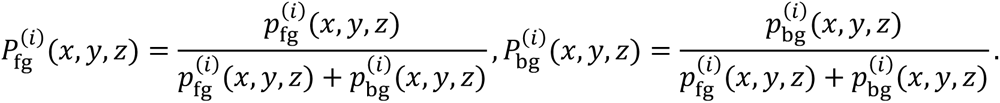

We modeled only these two classes.

*5. Aggregation across features to form a probability map*

To summarize evidence across features at each voxel, we computed per-feature foreground posteriors 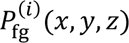 for all *i* = 1, …, *N*_feat_ and then formed an aggregated foreground confidence score:

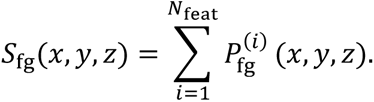

We stored *S*_fg_ as a 3D volume on the original (*x*′ *y*′ *z*) grid. This map provides a continuous-valued voxel-wise confidence measure; higher values indicate that a voxel’s feature intensities more consistently match the empirically learned foreground distributions across the feature set.

#### Segmentation using Probability Map

The 3D probability maps were subjected to Voronoi–Otsu labeling for object segmentation.

#### Spatio-temporal Tracing

We imported segmentation masks for each 3D time point from labeled TIFF stacks (one enumerated label per object). For each label, we extracted all voxel coordinates, computed the centroid as the mean voxel coordinate, and defined object volume as the voxel count. We tracked objects between consecutive time points (time reference, *tref*; time candidate, *tcan = tref + 1*).

We began with a *geometric assignment*. For each reference label (objects at *tref*), we identified candidate labels (objects at *tcand*) whose centroids lay within 3 voxels (Euclidean distance). When this criterion yielded an unambiguous nearest-neighbor match, we assigned the pair immediately and removed it from both the reference and candidate label lists, preventing entry into subsequent stages.

We matched the remaining objects by *features*. For each reference label, we restricted candidate labels to those within a fixed spatial window centered on the reference centroid (typically a 50 × 50 × 20 voxel box in x, y, z), limiting comparisons to spatially plausible pairs. For each time point, SpatialDINO produced 390 feature volumes. For every object, we computed a minimal bounding box, cropped the corresponding feature subvolume from the global 3D feature stack, and extracted the associated binary mask. We aligned each reference–candidate pair by shifting the candidate subvolume so that its mask centroid coincided with the reference centroid. We embedded both masks in a union volume large enough to contain them after alignment, enabling voxel-by-voxel comparison without rescaling or interpolation. For each aligned reference–candidate pair, we computed similarity directly on the voxel grids. We quantified binary similarity with the Dice coefficient. For each feature channel, we compared voxel-wise feature values using Pearson correlation and computed mean squared error (MSE) over all overlapping voxels. We based these metrics strictly on the feature values and evaluated all comparisons voxel-by-voxel.

For each reference label, we screened each associated candidate against minimum thresholds of 0.5 for Dice and 0.5 for Pearson correlation, enforcing similarity in volume, shape, and gradient-like feature structure. We retained only threshold-passing pairs for feature-wise selection. We then ran a feature-wise tournament. For each individual feature, we compared voxel-wise MSE across all retained candidate–reference pairs, selected the candidate with the lowest MSE as that feature’s best match, and awarded it one win. Thus, each feature contributed exactly one win to a single candidate.

After evaluating all 390 features, we summed wins for each candidate label associated with a given reference label and recorded the contributing feature set. We assigned, for that iteration, the candidate with the highest win total as the match for a reference label if it was the winner for only one reference and if its win total exceeded a minimum winning threshold. We decreased this threshold across iterations (340, 320, 300, 280, and 260 wins), removing assigned pairs from both reference and candidate lists after each iteration. Because each assignment changes the competitive set, we recomputed win counts from scratch after each match assignment and iterated until no additional pairs met the current threshold.

After the feature-threshold competition removed all high-confidence assignments, we linked remaining reference labels by a final *geometric proximity* rule. For each unassigned reference object, we selected the candidate label with the smallest 3D centroid distance among its candidate list. If this minimum distance lay within the allowed spatial search window, we assigned the pair as a residual match, retaining objects with weak feature support but clear geometric proximity. We recorded as unmatched any remaining reference object with no viable candidate within the search radius.

### 4D Data viewer

We developed *<u>Mirante4D,</u>* an interactive viewer for 4D (3D + time) fluorescence microscopy datasets; the code is openly available at https://github.com/kirchhausenlab/llsm_viewer/ and deployed as a static website at https://kirchhausenlab.github.io/llsm_viewer/. Although we designed Mirante4D for multi-channel fluorescence volumes, it also accepts other volumetric modalities (e.g., MRI), provided the data can be represented as one or more co-registered monochromatic channels sampled on a 3D grid as a single volume frame or over time. The site runs as a single-page application without a server-side backend, so all processing and rendering occur locally in the user’s browser, and no user data leave the local machine. The viewer accepts multi-channel volumetric image data, volumetric segmentation masks, and associated particle/track annotations. It provides both 2D slice-based views and 3D volume renderings, with GPU-accelerated visualization implemented in WebGL (Three.js with custom shaders).

To maintain interactive performance while constraining memory usage, we preprocess raw TIFF stacks into a browser-streamable “preprocessed dataset” format. During viewing, the application fetches volumes on demand from the preprocessed dataset rather than materializing entire time series in memory, enabling interactive inspection of long 4D recordings.

The viewer includes utilities for anisotropy correction, track-amplitude plotting, isosurface rendering, movie recording, and per-channel XY registration for colocalization analyses. It also includes an experimental virtual-reality mode for use with VR headsets.

### Hardware

The computer infrastructure supporting SpatialDINO training and GPU-based inference comprised three NVIDIA DGX A100 systems, each equipped with eight A100 40-GB GPUs interconnected by NVSwitch, providing 600 GB/s NVLink connectivity among all GPUs. Each DGX A100 node included two AMD EPYC 7742 64-core CPUs, 2 × 1 TB DDR4 system memory, two 1.92-TB M.2 NVMe SSDs for the operating system, and four 3.84-TB PCIe Gen4 NVMe SSDs for local storage. Networking included dual 10/25/40/100/200 GbE QSFP28 Ethernet ports via a ConnectX-6 adapter, eight Mellanox ConnectX-6 QSFP28 200-Gbps HDR InfiniBand/200GigE adapters, and dedicated management (RJ45) and BMC ports. Systems ran Ubuntu.

CPU-based postprocessing used a cluster of ∼800 CPU cores connected via 10 Gb/s links to laboratory workstations.

### Data Availability

All datasets and codebase are available upon request. For further information, please contact the lead author at.

## FIGURE AND VIDEO LEGENDS

**Figure S1.**
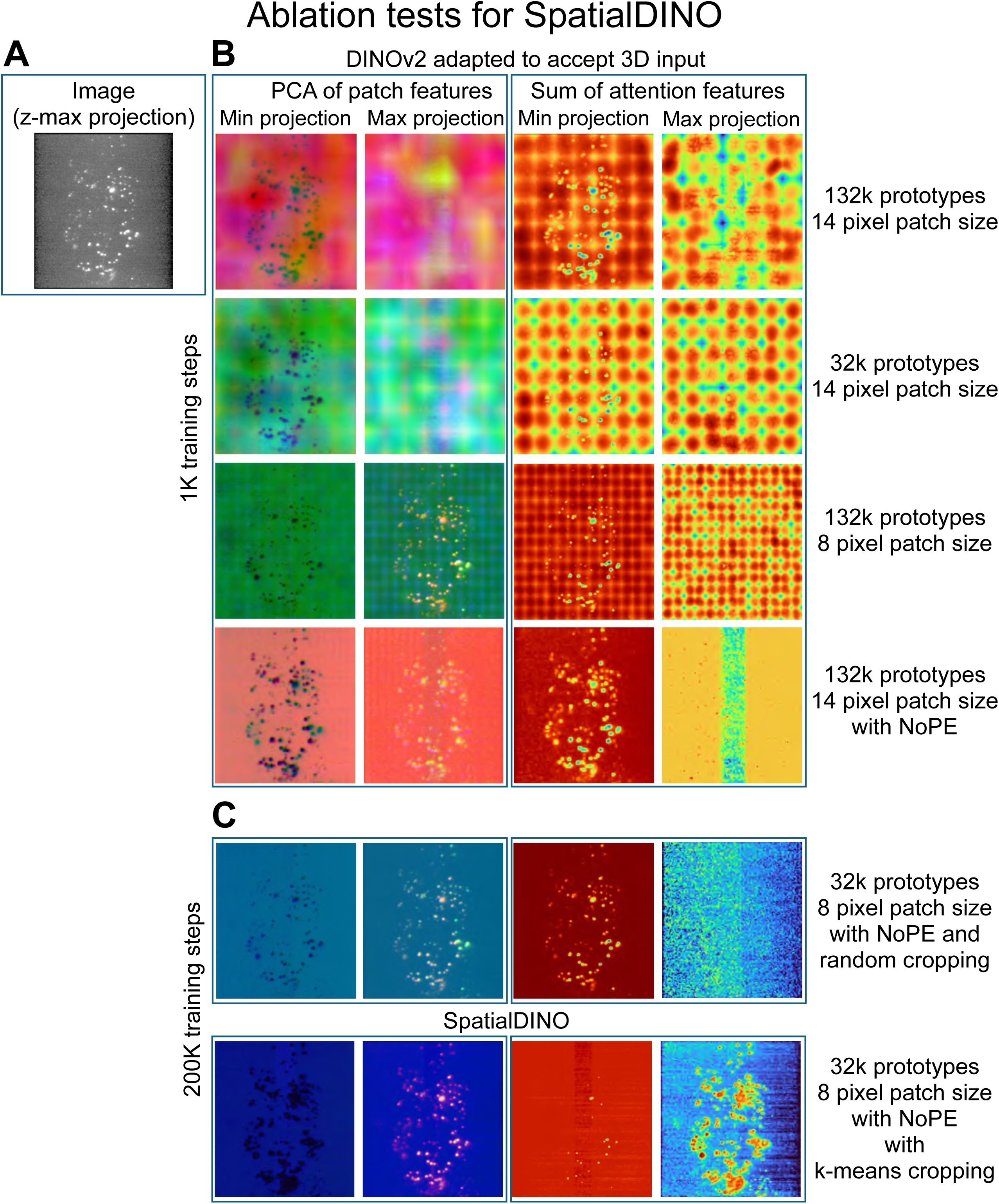
SpatialDINO ablations. Associated to Fig. 4A. **(A)** Max z-projection of the image in Fig. 4A corresponding to a 3D LLSM image that had not been used during training of endosomes and lysosomes labeled with internalized fluorescently labelled dextran. **(B)** Patch and attention features for early training, after 1K iterations. The baseline (1st row, top to bottom) corresponds to the original DINOv2, merely adapted to accept 3D inputs, and shows very strong grid and high-norm artifacts in the feature maps. The quality of feature maps barely changes after the four-fold reduction in number of prototypes (feature dimension of the projection heads), as shown in the 2nd row. In the 3rd row, the reduction in patch size from 14 to 8 voxels led to both finer features and reduced strength of the artifacts. Finally, only with the removal of positional encodings (known as NoPE, 4th row) were the features made free of artifacts. **(C)** Patch and attention features for late training, after 200K iterations. Top row shows the features for SpatialDINO without k-means cropping augmentation for global and local views. The bottom row shows the features for the final SpatialDINO configuration, with k-means cropping. In this case, the attention features were able to properly “attach” to foreground objects for their maximum and background noise for their minimum.

**Figure S2.**
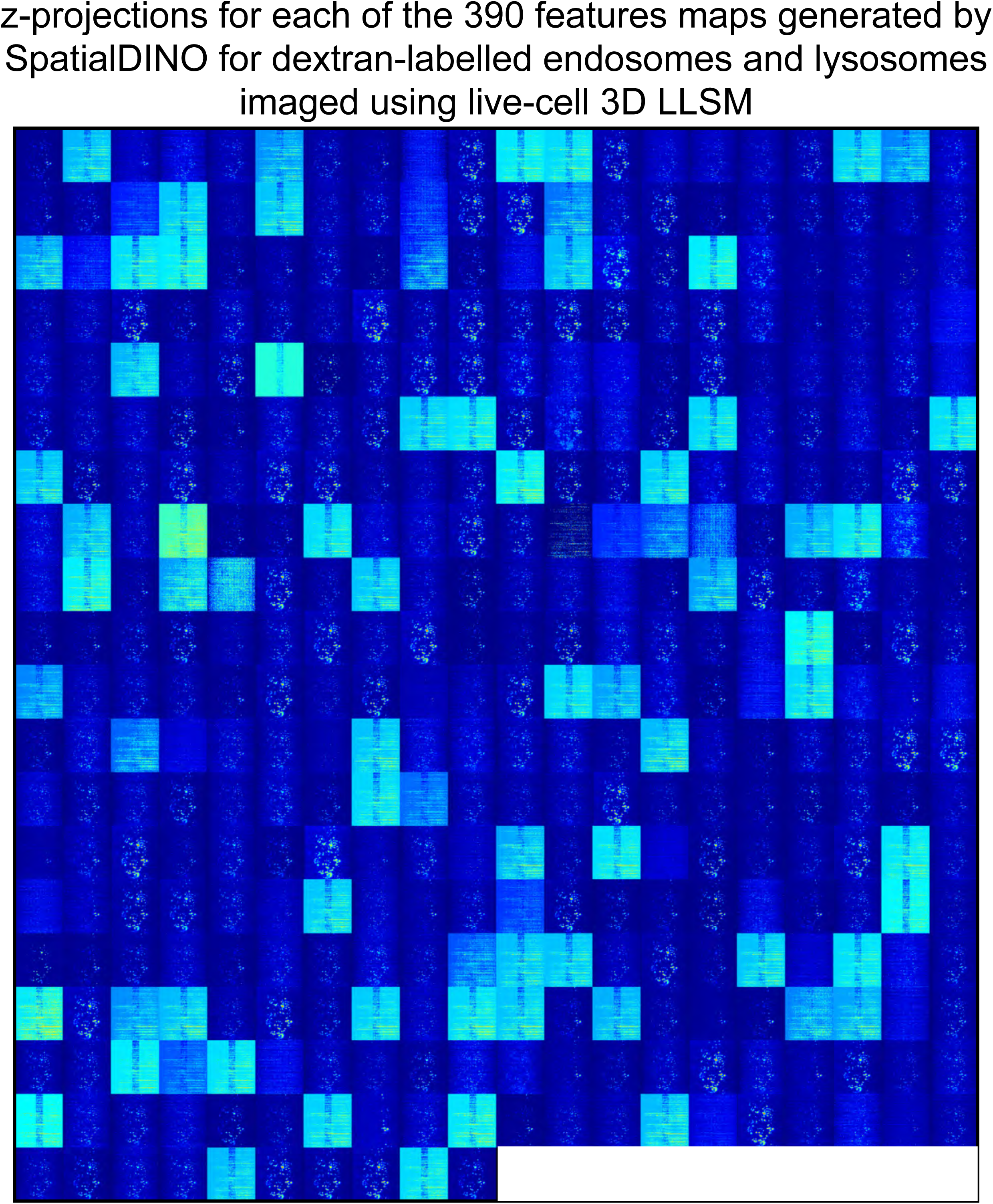
Normalized maximum-intensity projections of 390 SpatialDINO feature maps for dextran-labeled endosomes and lysosomes. Associated with Fig. 4A. Representative example showing voxelwise z–maximum-intensity projections for each of the 390 SpatialDINO feature maps inferred from a volume such as in Fig. 4A (Dextran-640–labeled endosomes and lysosomes imaged by live-cell 3D LLSM). Feature maps are arranged from feature 0 (upper left) to feature 389 (lower right) and normalized for display.

**Figure S3.**
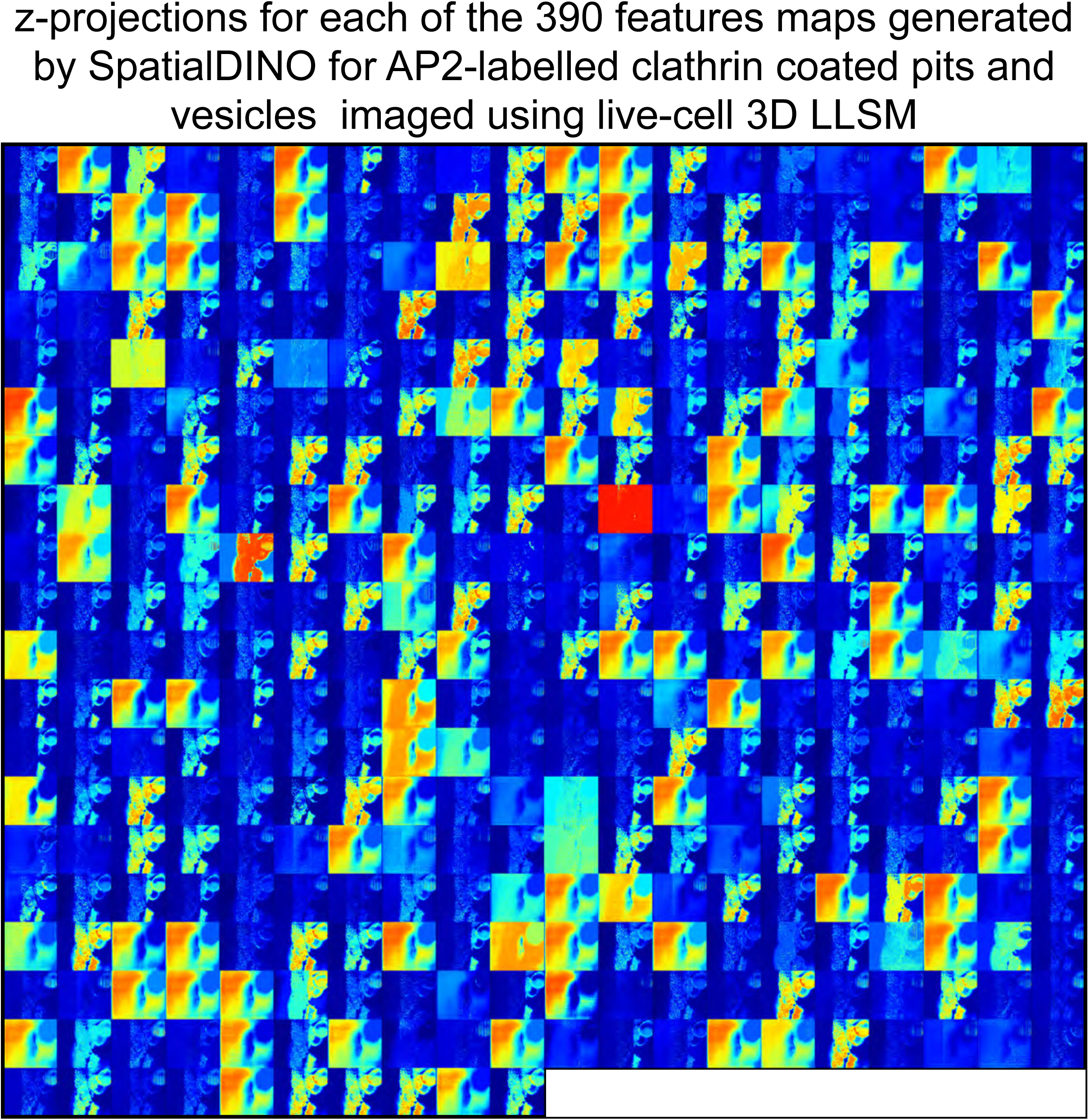
Normalized maximum-intensity projections of 390 SpatialDINO feature maps for AP2-marked clathrin-coated structures. Associated with Fig. 5A. Representative example showing voxelwise z–maximum-intensity projections for each of the 390 SpatialDINO feature maps inferred from a live-cell 3D LLSM volume of clathrin-coated pits and vesicles labeled with AP2 (as in Fig. 5A). Feature maps are arranged from feature 0 (upper left) to feature 389 (lower right) and normalized for display.

**Figure S4.**
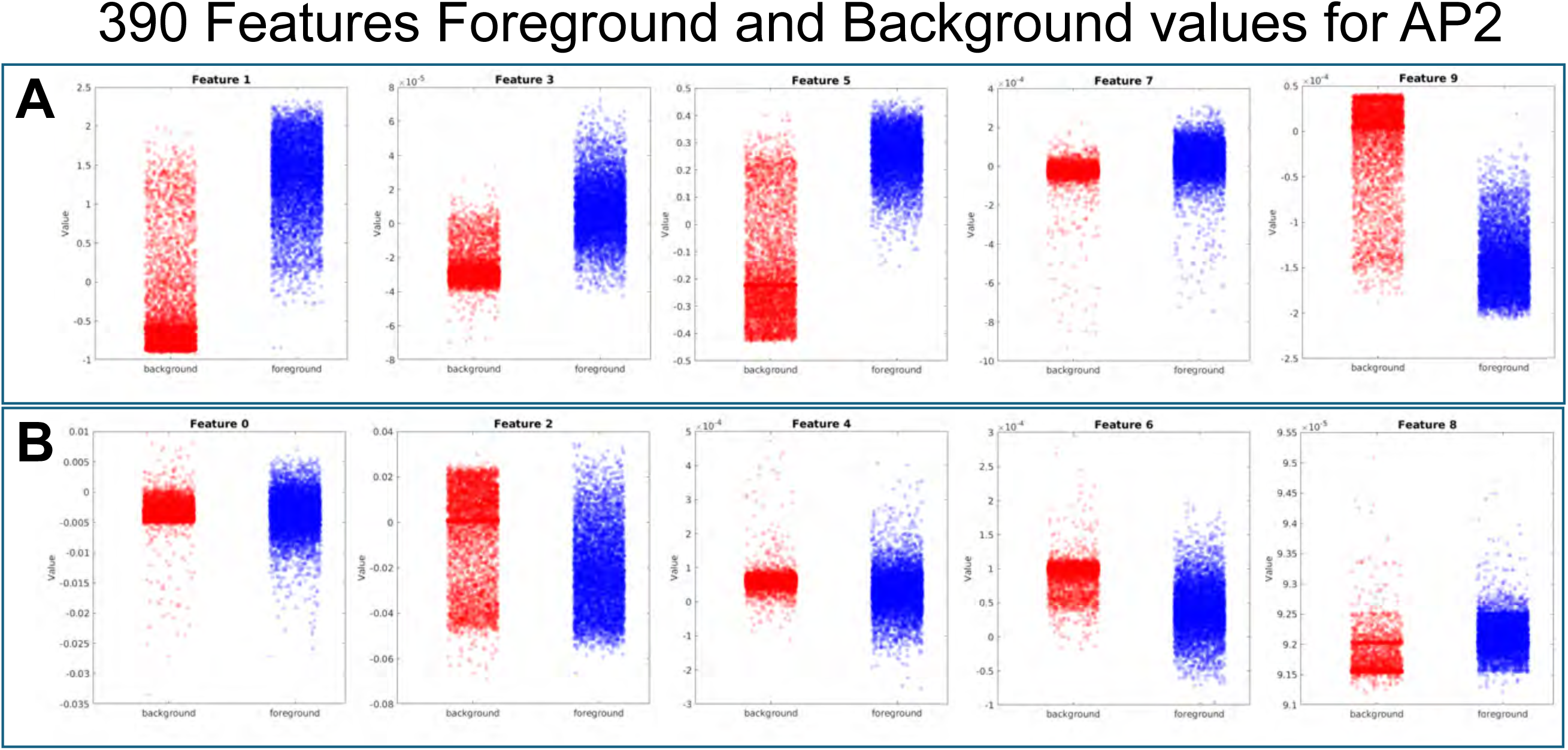
SpatialDINO feature maps derived from 3D CME-defined foreground and background voxels in AP2-labeled clathrin-coated structures. Associated with Fig. 5A. Voxelwise values for each of the 390 SpatialDINO features were computed from live-cell 3D LLSM volumes of AP2-labeled clathrin-coated pits and vesicles (as in Fig. 5A). Foreground (coated structures) and background (cytosol) voxels were defined by a 3D CME-based segmentation derived from the raw images. Plots show the distributions of feature values for foreground (red) and background (blue) voxels, illustrating feature-specific separation. **(A)** Selected representative SpatialDINO features showing detectable separation between foreground and background: the voxel-value distributions substantially overlap are distinguishable. **(B)** Selected representative SpatialDINO features not showing detectable separation between foreground and background: the voxel-value distributions substantially overlap and are not distinguishable.

**Figure S5.**
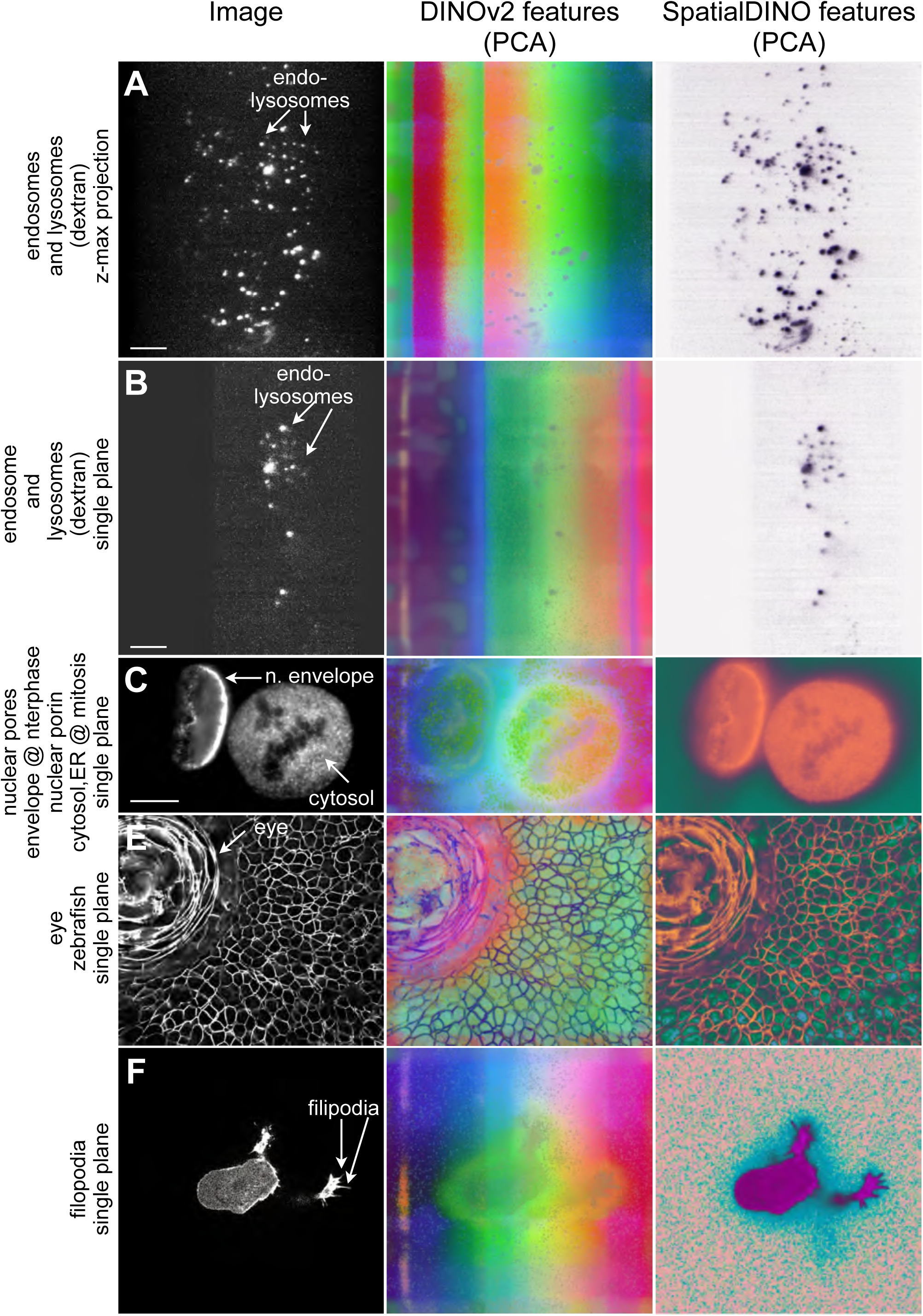
Comparison of DINOv2 and SpatialDINO inference on 2D sections from 3D fluorescence volumes (related to Fig. 4). For each example, we show (left to right) the raw image and an RGB rendering of the first three PCA components of the DINOv2 and SpatialDINO feature maps. We used the public DINOv2 ViT-L/14 checkpoint with registers and upsampled the input 5.25× to match SpatialDINO’s effective sampling (3× upsampling with patch sizes 14 versus 8). We applied DINOv2 to each 2D optical section independently (“2.5D” inference) and computed PCA over the full 3D volume. **(A)** Endosomes and lysosomes in SUM159 cells after 3 h uptake of fluorescent dextran (Dextran–Alexa647), imaged by live-cell 3D LLSM. Similar image types were included in SpatialDINO training. Z-max projections are shown; here DINOv2 was applied directly to the z-max projection. Scale bar, 10 µm. **(B)** Same as **(A)** but showing a single optical section instead of the z-max projection. **(C)** Halo–Nup133 in gene-edited SUM159 cells labeled with Janelia Fluor 549, imaged by live-cell 3D LLSM (acquired from (12)). This image type was not used for SpatialDINO training. A single optical section is shown. Scale bar, 10 µm. **(D)** Zebrafish eye and surrounding tissue with membranes labeled by an mStayGold fusion, imaged by live-cell oblique light-sheet microscopy and spatially deconvolved. This image type was not used for training. A single optical section is shown. Scale bar not available. **(E)** HBEC cell ectopically expressing tractin–GFP, imaged by two-photon Bessel-beam light-sheet microscopy followed by spatial deconvolution (14, 15, 45). This image type was not used for training. Scale bar not available.

**Video S1. SpatialDINO-based visualization of the cell surface and filopodia. Associated to Fig. 6**. Mirante4D volumetric rendering of an HBEC cell expressing tractin–GFP (45). The movie shows, in sequence, the deconvolved light-sheet volume, the SpatialDINO-derived probability map, and surface renderings of segmentation masks generated by u-Segment 3D (45) (white) and by local Otsu thresholding of the SpatialDINO probability map (blue).

**Video S2. SpatialDINO-based visualization of breast MRI. Associated to Fig. 7**.

Mirante4D rendition of the 3D MRI volume for the left and right breasts (DUKE: Breast_MRI_141). MRI volume is shown first and then overlaid with binary segmentations derived from SpatialDINO-based probability maps generated using lesion foreground–background labels from the ipsilateral breast highlighting left-breast– derived mask (green) and right-breast–derived mask (red).

**Video S3. SpatialDINO-based tracking of simulated endosomes. Associated to Fig. 8B**.

Mirante4D volumetric visualization of a simulated 3D fluorescence endosome movie. The movie first shows the simulated endosomes alone and then overlays SpatialDINO-derived trajectories, displayed initially as the current position with a two-frame history and subsequently with the full track history.

**Video S4. SpatialDINO instance segmentation of endosomes and lysosomes. Associated to Fig. 8C**. Mirante4D volumetric visualization of an SVG-A cell imaged by live-cell 3D LLSM containing dextran-labeled endosomes and lysosomes. The raw volume is shown first, followed by an overlay with the SpatialDINO instance-segmentation mask, with a distinct pseudo color assigned to each object.

## ACKNOWLEDGMENTS

We thank members of the Kirchhausen laboratory for advice and encouragement; Justin O’Connor for help maintaining the computing infrastructure; and Gokul Upadhyayula and Eric Betzig for providing the zebrafish dataset imaged by Amir Hay using oblique light-sheet microscopy with reagents prepared by Dave Matus. This work was supported in part by a National Institute of General Medical Sciences Maximizing Investigators’ Research Award R25 GM130386 to Kirchhausen, the NNF Center of Optimized Oligo Escape and Control of Disease grant to T.K. and N.H., and by discretionary funds to T.K. G. S. was funded in part by a National Institute of Health grant R01CA272484 to T. Kirchhausen and Stephen Blacklow.

Acquisition of the computing hardware (DGX GPU systems, CPU clusters, high-speed storage, archival servers, and workstations) was supported by a Massachusetts Life Sciences Center Pilot grant to T. Kirchhausen. and by an equipment supplement to GM130386 to T. Kirchhausen. Construction of the server room was supported by generous support from the PCMM Program at Boston Children’s Hospital.

**The authors declare no competing financial interests.**

## Author contributions

Alex Lavaee conceived the approach for adapting DINOv2 to native 3D data and helped define the strategy underlying SpatialDINO. Alex Lavaee and Arkash Jain developed, tested, and implemented SpatialDINO. Gustavo Scanavachi Moreira Campos and Jose Inacio Costa-Filho developed the pipelines for instance segmentation and trajectory reconstruction from SpatialDINO-inferred feature volumes. Jose Inacio Costa-Filho developed and implemented the 4D viewer, Mirante4D. Adam Ingemansson performed the initial quantitative analysis of SpatialDINO segmentation on clathrin-coated pits and vesicles. Tom Kirchhausen introduced the problem, defined the overall study logic, drafted the manuscript, and finalized it in consultation with all authors.

---

**Table SI.**
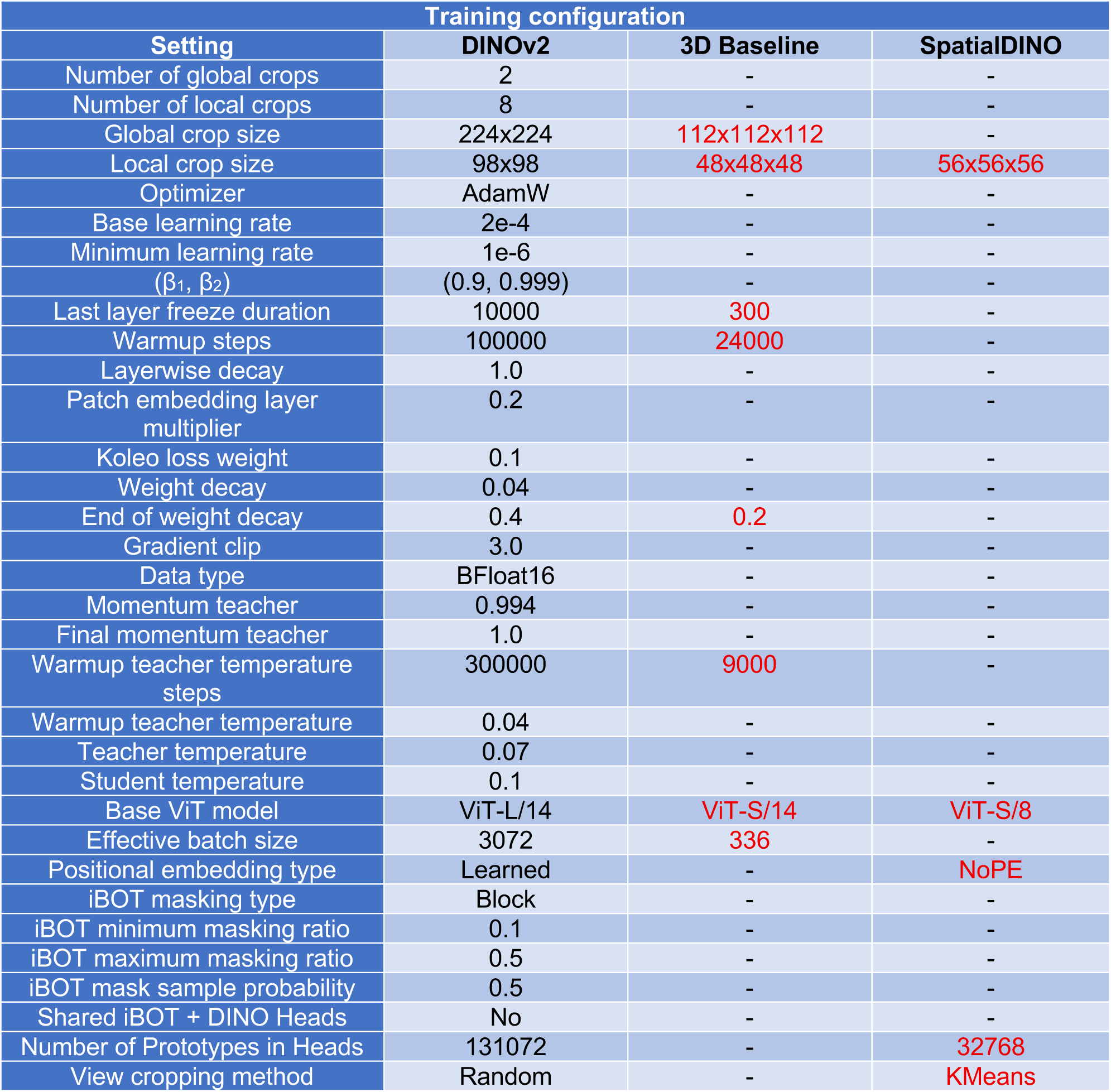

